# Inhibition of IL2Rß-dependent STAT5 activity supports T-cell stemness and augments antitumor efficacy of CD8^+^ T cells by preventing T-cell exhaustion

**DOI:** 10.1101/2025.05.14.651875

**Authors:** M Shourian, B Bourdin, N El-Hachem, C Ruisseaux, A Vallée, H Roméro, EO Kwarteng, E Haddad, VP Lavallée, JC Beltra, H Decaluwe

## Abstract

CD8^+^ T-cell exhaustion is a leading cause of adoptive cell therapy (ACT) failure. In contrast, maintaining a stem-like state correlates with better expansion, persistence, and anti-tumor activity of infused T-cell products. IL-2 is extensively used in ACT protocols given its ability to expand T-cell populations. Yet, IL-2 drives more differentiated and exhausted states, diminishing the quality of T-cell products. Understanding how cytokines of the IL2R family drive T-cell differentiation is essential to ultimately design optimal ACT protocols, safeguarding stem-like programs while ensuring sufficient T-cell expansion. Here, we show that cytokine signaling through IL2Rβ supports more differentiated exhausted T cells in chronic lymphocytic choriomeningitis infection. Similarly, high levels of IL-2 and IL-15 *in vitro* foster heightened differentiation and exhaustion of cells for adoptive cell therapy. In contrast, absence of IL2Rβ in vivo or transient inhibition of Janus kinase 3 (JAK3) or signal transducer and activator of transcription 5 (STAT5) *in vitro* favors features of T-cell stemness. Transcriptional analyses of *in vitro* expanded T cells further reveal that inhibition of STAT5 sustains a stemness program, which correlates with better antitumor activity in a mouse melanoma model. When applied to a human CAR T expansion model, inhibition of STAT5 supports memory progenitor differentiation and limit inhibitory receptor expression. These results demonstrate that continuous exposure to high levels of cytokines, such as IL-2 and IL-15, constrain CD8^+^ T cells towards more advanced states of exhaustion. In contrast, limiting cytokine signaling using specific kinase inhibitors preserves stem-like T-cell programs and enhance the quality of ACT products.

**One Sentence Summary:** Sustained IL-2/IL-15 signaling drives CD8^+^ T-cell exhaustion while JAK3/STAT5 inhibition preserves stemness, boosting adoptive cell therapy efficacy.

## INTRODUCTION

Adoptive cell therapy (ACT) can elicit profound therapeutic responses, including durable remission in patients with advanced malignancies (*1*). Despite these clinical successes, most patients undergoing ACT do not experience long-term benefits (*1, 2*). Hence, there are considerable interests in optimizing ACT approaches to achieve better response rates in patients with cancer (*3*). Hallmarks for highly effective ACT includes the maintenance of stem-like T cells in ACT products that typically conserve self-renewal potential and show features of enhanced proliferation, *in vivo* persistence, and superior anti-tumor activity (*4*). However, despite being central for therapy success, these stem-like cells typically form a minor proportion of ACT products (*5, 6*). Strategies to optimize ACT protocols are needed to increase the yield and quality of stem-like T cells while coping with sufficient numerical expansion of T-cell products for downstream therapeutic applications (*7*).

Cytokines of the common gamma (γ_c_) chain family direct the expansion and differentiation of CD8^+^ T cells during infection and cancer. These cytokines (interleukin (IL)-2, −4, −7, −9, −21) signal through inherent receptors and activate specific combinations of signaling molecules to exert their non-redundant functions on T-cell differentiation and cell-fate commitment. The affinity and strength of binding and specificity of signaling kinases dictate CD8^+^ T-cell programs and outcome (*8–10*). In ACT protocols, IL-2 is primarily used to support T-cell expansion, but IL-2-derived signals also trigger terminal differentiation at the expense of stemness (*8, 11, 12*). To overcome this major limitation, alternate combinations of γ_c_ cytokines (e.g. IL-7, −15, −21) have been employed to expand T-cell products, with improvements on subsequent antitumor activity (*13–15*). Although these findings are promising, our understanding of how cytokine signaling influence T-cell fate commitments remains limited. Few studies have explored the potential of cytokine kinase inhibitors to direct cells towards increased stemness and reduced exhaustion. Here, we report on the capacity to safeguard ACT stem-like states and antitumor functions by specifically targeting the IL2Rβ-dependent signaling cascade, common to both IL-2 and IL-15, with small molecule inhibitors of <u>ja</u>nus <u>k</u>inase 3 (JAK3) and <u>s</u>ignal transducer and <u>a</u>ctivator of transcription 5 (STAT5).

## RESULTS

### Lack of IL2Rβ-dependent signals during chronic lymphocytic choriomeningitis virus clone 13 infection maintains exhausted CD8^+^ T cells in a progenitor state

IL-2 and IL-15 cytokines have distinct but also redundant functions in CD8^+^ T-cell immunity. Both cytokines share common receptor chains (IL2Rβ, IL2Rγ) and activate similar signaling molecules (e.g. JAK3, STAT5) in T cells (*16, 17*). To interrogate the importance of IL-2/IL-15-dependent signals on CD8^+^ T-cell differentiation fates in the context of exhaustion, we took advantage of the murine lymphocytic choriomeningitis virus (LCMV) clone 13 (Cl.13) model. This model causes a chronic infection in mice and exhaustion of virus-specific CD8^+^ T cells by ∼1 month of infection (*18*). Using this model, we previously found that expression of IL2Rβ (CD122; common to IL-2 and IL-15) on antigen (Ag)-specific CD8^+^ T cells positively correlated with features of T-cell exhaustion (e.g. IR expression) and extended these observations to virus-specific CD8^+^ T cells in patients with chronic hepatitis C (*18*). To interrogate how IL2Rβ expression influences T-cell commitments towards exhaustion, we used the same model and sorted CD8^+^ T cells specific for the gp33-41 epitope of the LCMV virus based on their level of expression of IL2Rβ at day 21 post infection (p.i.) with LCMV C.13. Bulk RNA sequencing was then performed and genes from cells with low (IL2Rβ^lo^) or high (IL2Rβ^hi^) IL2Rβ expression compared (n=3 mice per group). At a false discovery rate (FDR) of 5% and using a log2 fold change (FC) ≥ 0.5, 1,293 differentially expressed (DE) genes were upregulated and 1,408 DE genes were downregulated in IL2Rβ^hi^ compared to IL2Rβ^lo^ cells (**Fig. 1A**, **Table S1**). Comparative gene set enrichment analyses (GSEA) further showed a striking enrichment for terminally exhausted signature genes (when compared to Slamf6^-^ Tim3^+^ CD8^+^ T cells from day 28 pi)(*19*) among DE genes of cells expressing high levels of IL2Rβ, while progenitor exhausted signature genes (from Slamf6^+^ Tim3^-^ CD8^+^ T cells from day 28 pi)(*19*) were downregulated (**Fig. S1A-B**). The upregulated genes in IL2Rβ^hi^ cells include those encoding inhibitory receptors such as *Pdcd1* (PD1), *CD244a* (2B4), *CD160*, *Lag-3*, *Tigit*, *Havcr2* (TIM-3), *CD101*, *CD38*, and *Entpd1* (CD39), CD200R2, and CD200R4; costimulatory molecules like *Tnfrsf9* (CD137 or 4-1BB) and *Tnfrsf7* (CD27); cytotoxic proteases such as *Gzma*, *Gzmb*; and transcription factors including *Tox*, *Prdm1* (Blimp), *Eomes*, *Ikzf2,* and *Batf* (**Fig. 1A**, **Table S1**). Conversely, genes associated with T-cell stemness were downregulated in IL2Rβ^hi^ gp33-41-specific cells. These include surface receptors such as *Il7r*, *Ccr7*, *Slamf6* (Ly108), *Sell* (CD62L) and *S1pr1*; the effector molecule *TNF*; and transcription factors such as *Id3*, *Tcf7* (TCF-1), *Lef1*, *Satb1* and *Klf2* (**Fig. 1A**, **Table S1**). These observations suggest a role for sustained IL2Rβ-dependent signals in fostering more differentiated exhausted states at the expense of T-cell stemness in context of persistent antigenic stimulation.

**Fig. 1.**
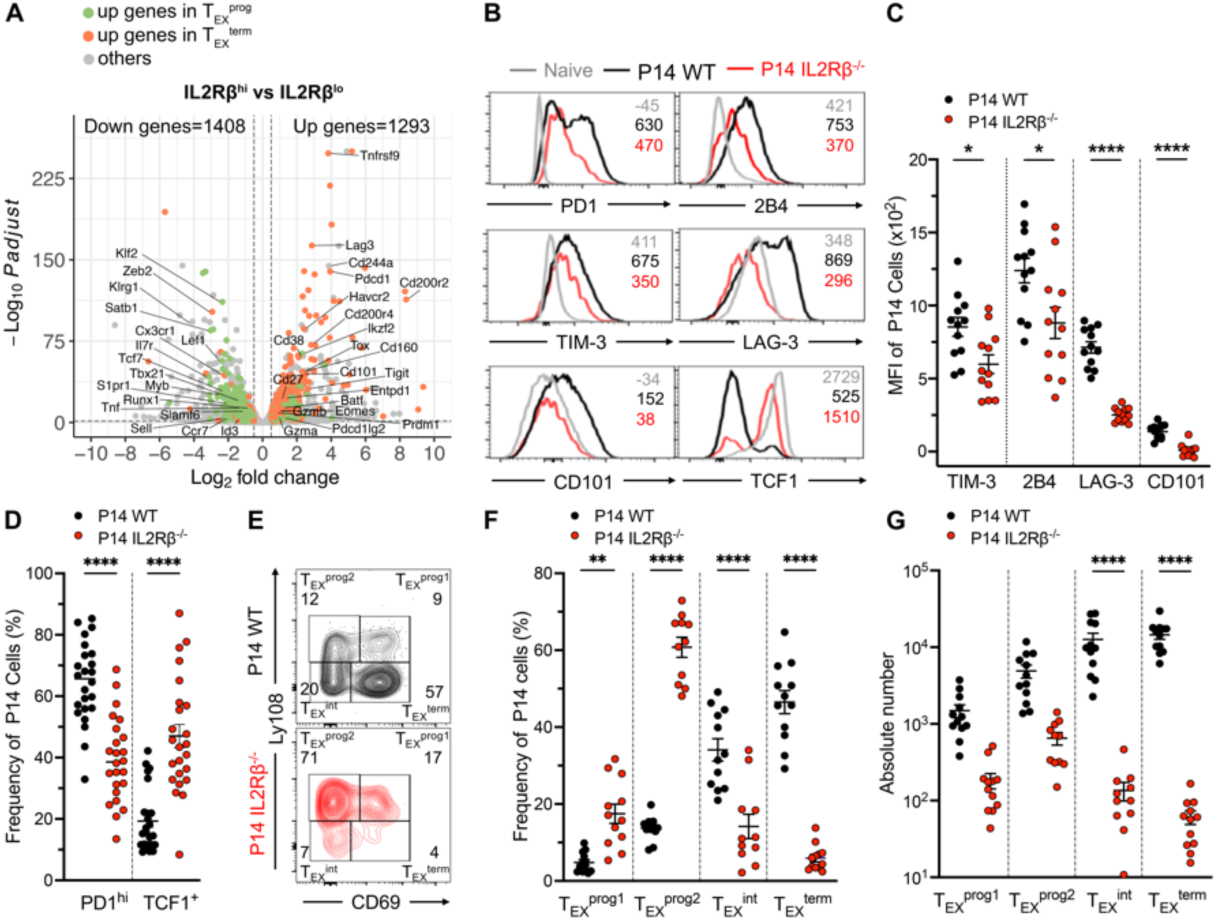
The absence of IL-2Rβ signaling sustains CD8^+^ T cells in a progenitor exhausted state during LCMV Cl.13 infection. (**A**) Volcano plot illustrating DE genes between IL2Rβ^hi^ and IL2Rβ^lo^ isolated CD8^+^ Tet^+^ CD11a^+^ T cells at day 21 p.i. Genes significantly upregulated in T_EX_^prog^ (Slamf6^+^Tim-3^-^, green) and in T_EX_^term^ (Slamf6^+^Tim-3^+^, orange) cells according to Miller *et al.* (GSE123235) (*19*) or not (grey) are depicted (log₂ FC ≥ 0.5 and FDR < 5%; n= 3 biological replicates per group). (**B-G**) Spleen cells analyzed at day 28 p.i. of adoptive co-transferred of P14^+^ IL2Rβ^+/+^ (P14 WT, black line or circles) and P14^+^ IL2Rβ^-/-^ (red line or circles) cells into B6 mice. Histograms (**B**) and scatter plots (**C**) of the MFI of indicated surface markers. (**D**) Scatter plots of the frequency of PD1^hi^ and TCF1^+^ cell populations. (**E**) Flow cytometry plots for Ly108 and CD69 co-expression. Scatter plots of the frequency (**F**), and absolute number (**G**) of T_EX_^Prog1^ (CD69^+^Ly108^+^), T_EX_^Prog2^ (CD69^-^Ly108^+^), T_EX_^int^ (CD69^-^Ly108^-^), and T_EX_^term^ (CD69^+^Ly108^-^) cell populations. Data are representative of (C, F and G) or combined (D) from two independent experiments (total of 24 mice). Error bars indicate means ± SEM. *P < 0.05; **P < 0.01; ***P < 0.001; ****P < 0.000, using Mann-Whitney (C, D) and one-way ANOVA (F, G).

To specifically address whether IL2Rβ expression on CD8^+^ T cells supports the transition from progenitor to more differentiated exhausted states *in vivo*, we co-transferred 5 × 10^3^ congenically distinct populations of wild-type (WT, CD45.1/CD45.2) and IL2Rβ-deficient (IL2Rβ^-/-^, CD45.2) CD8^+^ T cells transgenic (Tg) for the gp33-41 epitope of LCMV (P14 mice) into C57BL/6 (B6, CD45.1) recipient mice one day prior infection with LCMV Cl.13 (**Fig. S1C**). At d28p.i., when exhaustion was fully established (*20–22*), donor cells isolated from the spleen were analyzed by flow cytometry. We observed a greater accumulation of WT over IL2Rβ^-/-^ P14 cells (∼37-fold) (**Fig. S1D**). Phenotypically, P14 R IL2Rβ^-/-^ cells displayed significantly decreased expression of all IRs tested – i.e. 2B4, TIM-3, Lag3, CD101 – compared to P14 WT, and a reduced frequency of PD1^hi^ cells (**Fig. 1B-D**). At this time-point during a chronic LCMV infection, distinct but developmentally related subsets of exhausted T cells (T_EX_) have been identified. These include TCF1-expressing progenitors (Ly108^+^ CD69^+^/ T_EX_^prog1^ and Ly108^+^ CD69^-^/ T_EX_^prog2^) that continuously repopulate more differentiated populations including an intermediate or “effector-like” subset (Ly108^-^ CD69^-^ or Ly108^-^ CX3CR1^+^ / T_EX_^int^), and a terminally exhausted population (Ly108^-^ CD69^+^ or Ly108^-^CX3CR1^-^/ T_EX_^term^) that both loose TCF1 expression (*22, 23*). In the absence of the IL2Rβ-chain, increased expression of TCF1 suggested a developmental bias toward progenitor/stem-like exhausted cells (**Fig. 1B, D**). Consistently, the proportion of distinct T_EX_ subsets was redistributed towards a higher frequency of Ly108-expressing T_EX_ progenitors in IL2Rb-deficient cells compared to P14 WT (both Ly108^+^ CD69^-^ T_EX_^prog2^ and Ly108^+^ CX3CR1^-^ stem-like cells) (**Fig 1E-F, S1E-F**). In contrast, the frequency of T_EX_^int^ (defined either as Ly108^-^ CD69^-^ or CX3CR1^+^) and T_EX_^term^ (Ly108^-^CD69^+^ or Ly108^-^CX3CR1^-^) cells were reduced in P14 Rβ^-/-^ compared to their WT counterpart. Absolute numbers reflected this developmental bias with dramatic reductions of T_EX_^int^ and T_EX_^term^ cells in the absence of IL-2 and IL-15 signals while the number of T_EX_ progenitor cells was less affected (**Fig. 1G, S1G.)** These findings point to IL2Rβ-dependent signals as a key axis fostering more differentiated T_EX_-cell states at the expense of a progenitor or stem-like program during chronic antigenic stimulation.

### Sustained IL-2 and IL-15 stimulation supports exhausted T cell differentiation *in vitro*

Next, we aimed to investigate whether IL2Rβ-dependent signals influence features of T-cell exhaustion *in vitro* and if an *in vitro* approach could help dissect how IL2Rβ-signals dictate T-cell fates. Based on our prior work (*18*), we developed an *in vitro* system to evaluate the effects of continuous IL-2 and IL-15 stimulation in context of transient or continuous TCR stimulation – as we would observe in contexts of acute or chronic infection respectively – on T-cell differentiation. Naïve P14 CD8^+^ T cells were stimulated with gp33 peptide-loaded dendritic cells (100 ng/mL). On days 2, 4, and 6, CD8^+^ T cells were normalized and expanded in the presence or absence of gp33 peptide (1 ng/mL) and graded levels of IL-2 or IL-15 (**Fig 2A**). Consistent with prior observations (*18*), both IL-2 and IL-15 had pronounced impact on the induction of 2B4, CD160, and to a lesser extent on LAG-3 and TIM-3, in a dose-dependent manner (**Fig 2B**). While continuous TCR stimulation supported elevated expression of these inhibitory receptors, 2B4 and TIM-3 were strictly regulated by cytokine signals. In contrast, we observed gradual loss of stem-like features (e.g. CD62L, Ly108, TCF-1) with heightened levels of both IL-2 and IL-15, particularly in context of transient peptide priming (**Fig 2C-D**). This suggested that constant IL-2 and IL-15 stimulation, when provided at high levels, prompts cells towards more differentiated T-cell fates.

**Fig. 2.**
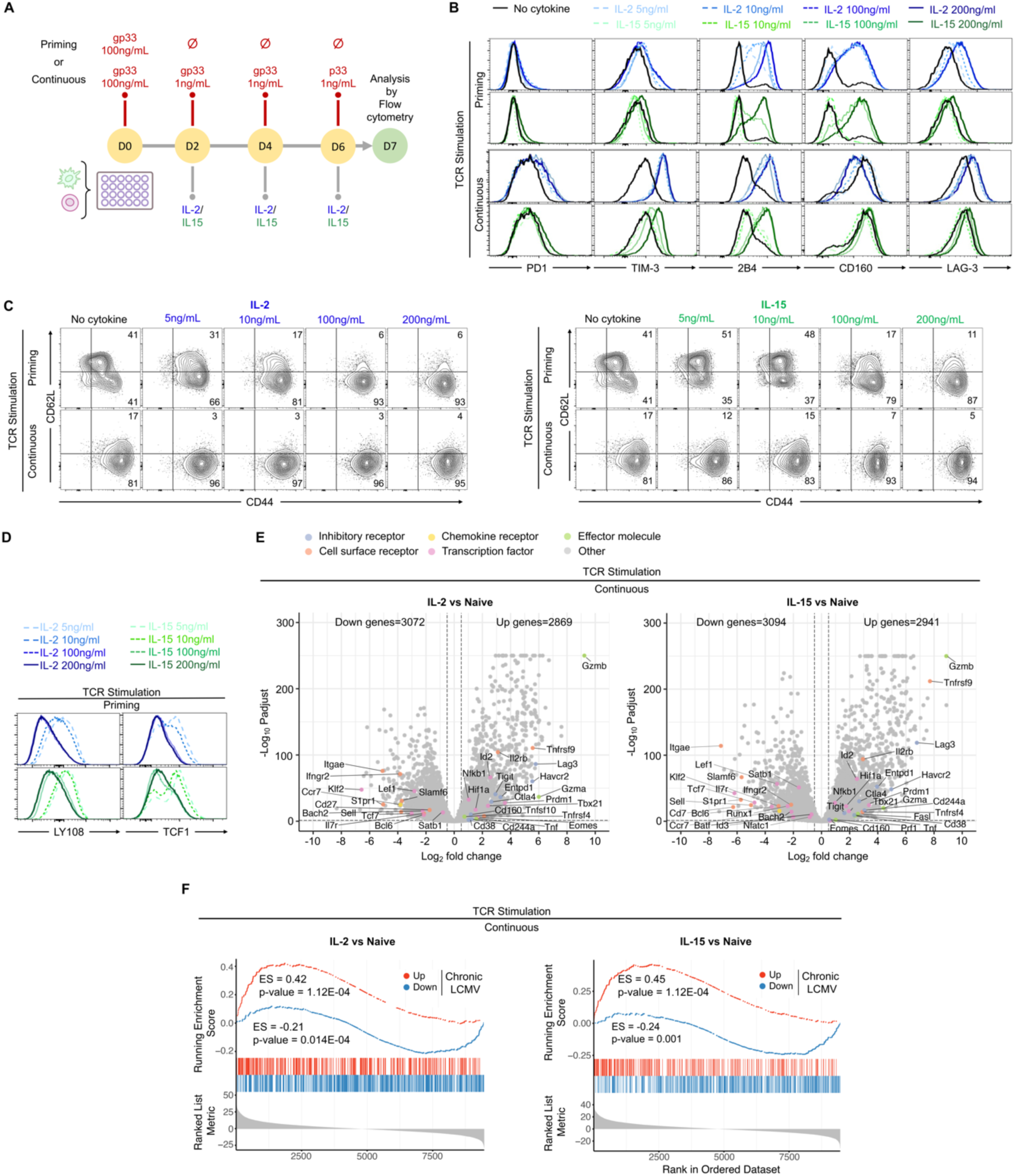
IL-2 and IL-15 enhance the differentiation of exhausted T cells *in vitro*. **(A)** Schematic of the experimental design for *in vitro* experiments. P14^+^ CD8^+^ T cells were co-cultured with gp33-pulsed DCs (100 ng/mL). On days 2, 4, and 6, P14^+^ cells were normalized and cultured without (priming) or with (continuous) gp33 peptide alone (1 ng/mL), or in combination with varying concentrations of IL-2 (blue) or IL-15 (green) (5, 10, 100, 200 ng/mL). Cells were analyzed at day +7. (**B**) Histograms of indicated IRs. **(C)** Contour plots for CD44/CD62L co-expression. (**D**) Histograms of the MFI of Ly108 and TCF1. (**E**) Volcano plots of the DE genes in continuously stimulated CD8^+^ T cells and IL-2 (left, 100 ng/mL) or IL-15 (right, 200 ng/mL) compared to naive CD8^+^ T cells (n= 3 replicates per group). Up- and down-regulated genes associated with T-cell exhaustion are labeled and categorized into pathways. Significance: log₂ FC ≥ 0.5 and FDR < 5% (**F**) Pre-ranked GSEA of DE gene profiles comparing IL-2- or IL-15-stimulated CD8^+^ T cells to naive CD8^+^ T cells against a T-cell exhaustion gene signature derived from Bengsch *et al.* (*24*) (upregulated genes: red line; downregulated genes: blue line). ES: enrichment score. Data are representative of three independent experiments (B-D).

To more precisely evaluate if our *in vitro* approach recapitulates features of T-cell exhaustion, CD8^+^ T cells stimulated with high levels of cytokines and continuous TCR stimulation were collected at the end of the culture for bulk RNA sequencing and compared to naïve CD8^+^ T cells (n=3 independent cultures per group). At a FDR of 5% and using a log2 FC ≥ 0.5, we found that 3,072 genes were upregulated and 2,869 genes were downregulated in IL-2 stimulated cells, while 3,094 genes were upregulated and 2,941 genes were downregulated in IL-15 stimulated cells, compared to naïve cells (**Fig 2E, Table S2-S3**). Continuous cytokines and TCR stimulation led to elevated expression of transcripts associated with terminally exhausted cells, including high levels of inhibitory receptors such as Lag-3, Havcr2 (TIM-3), Ctla4, Entpd1 (CD39), CD38, Tigit, CD244a (2B4), and CD160. Additionally, these cells showed increased levels of the transcription factors Prdm1 (BLIMP-1), Eomes, Hif1a, Nfkb1, and Id2, the cytolytic molecules Gzma and Gzmb, and the cell surface receptors Tnfrsf9 (CD137), Tnfrsf4 (CD134), and IL2Rβ. In contrast, the expression of transcription factors (i.e. *Satb1*, *Bach2*, *Tcf7* [TCF-1]*, Lef, Klf2*) and chemokines or cell surface proteins related to T-cell stemness (i.e. *IL-7r, Ccr7, S1pr1, Itgae, Sell* [CD62L)*, Slamf6* [Ly108]) were diminished in IL-2- or IL-15-stimulated CD8^+^ T cells. Notably, differential gene signatures observed in CD8^+^ T cells cultured with IL-2 or IL-15 and continuous TCR stimulation closely resembled the gene signatures of T_EX_ cells obtained from chronically-infected mice (*24*) (**Fig 2F**). Collectively, these results indicate that sustained TCR and IL-2 or IL-15 stimulation promotes the differentiation of T_EX_ cells *in vitro*, establishing our *in vitro* model as an effective system for investigating the cellular and molecular mechanisms underlying exhaustion.

### Inhibition of STAT5-signaling downstream of IL-2 and IL-15 enhances T-cell stemness

We next interrogated the signals downstream of IL-2 and IL-15 dictating CD8^+^ T_EX_ cell differentiation. IL-2 and IL-15 trigger various signaling pathways in CD8^+^ T cells, including the AKT/mTOR, ERK and JAK/STAT pathways (with preferential activation of STAT5 over STAT1 and STAT3) (*16, 25*). Activation of these pathways, however, might be deregulated in T_EX_ compared to conventional T_EFF_ or T_MEM_ cells with potential restrictions in AKT/mTOR and ERK activation in the former population (*26*). Leveraging the LCMV Cl.13 model, we probed which signaling pathway was fostering T_EX_ exit from the stem-like progenitor state to the terminally exhausted state. To do so, we isolated spleens from mice infected with LCMV Cl.13 at days 8 and 30 p.i. and assessed the downstream activation of key signaling molecules in P14 CD8^+^ T cells after *ex vivo* restimulation with IL-2 or IL-15. Naïve CD8^+^ T cells were also included as control. Consistent with prior work (*27, 28*), neither IL-2 nor IL-15 induced substantial activation of signaling molecules related to AKT/mTOR or ERK pathways (i.e. p-AKT, pERK, p-Foxo3a, pS6) in T_EX_ cells (**Fig 3A**). Induction of STAT1 and STAT3 phosphorylation was also not detected. In contrast, we observed substantial induction of phospho-STAT5 (pSTAT5) in the early phase of a LCMV Cl.13 infection in response to IL-2 and IL-15 (day 8 p.i) and the late phase of infection in response to IL-15 (day 28 p.i) (**Fig 3A**). These observations suggested a preferential role for the transcription factor STAT5 in acting downstream of IL-2 and IL-15-signals in T_EX_ cells *in vivo*. We next interrogated whether manipulating STAT5 activity (or JAK3 – its upstream kinase) would impact the developmental trajectory of CD8^+^ T cells in response to IL-2 or IL-15 cytokines *in vitro*. Using our exhaustion *in vitro* model described above, at doses of IL-2 (100 ng/mL) and IL-15 (200 ng/mL) which generated comparable T_EX_ phenotypes at the protein and transcriptional levels (**Fig 2**), small molecules inhibitors of STAT5 (STAT5i, **Fig 3B**) or JAK3 (JAK3i, tofacitinib – **Fig S3**) were added on days 4 and 6. Blocking STAT5 (**Fig 3C**) or JAK3 activity (**Fig S3B**) limited the induction of cytokine-inducible IRs – i.e. 2B4, TIM-3 and CD160. As for cytokines, the impact of STAT5 inhibition on IR expression was dose-dependent (**Fig S2A**). STAT5 inhibition also reinforced the expression of stem-like features in P14 CD8^+^ T cells including expression of CD62L, Ly108 and the canonical TF Tcf1 (**Fig 3D**, **S2B**). In fact, the addition of STAT5i on day 4 of the culture supported the development of cells presenting a progenitor-like state (Ly108^+^CD69^+^) and co-expressing markers of central memory T cells (T_CM_; CD62L^+^CD44^+^) in a dose-dependent manner (**Fig 3E-F, S2C-E**). STAT5i also strikingly limited the accumulation of terminally exhausted T_EX_^term^ (Ly108^-^CD69^+^) and effector/effector-like T cells (CD62L^-^CD44^+^ T_EFF_; Ly108^-^CD69^-^ T_EX_^int^), in sharp contrast with the IL-2 and IL-15 conditions (**Fig 3E-F, S2C-E**). JAK3 inhibition using either JAK3i or tofacitinib had a comparable effect on CD8^+^ T cell differentiation, with a greater impact of tofacitinib over JAK3i on the preservation of progenitor T-cells (**Fig S3D-G**). Consistent with the results obtained in the absence of IL2Rβ expression on CD8^+^ T cells *in vivo*, blocking JAK3-STAT5 signaling *in vitro* limited cell expansion at day 7 (by 1.6 to 1.7-fold) (**Fig S2F**). This correlated with reduced proliferation of CD8^+^ T cells in the presence of STAT5i (**Fig 3G, S2G**). However, the number of stem-like (Ly108-expressing) P14 cells emerging at the end of this *in vitro* culture was superior in conditions where STAT5i was added with striking increase in T_EX_^prog1^ and T_EX_^prog2^ (**Fig 3H**). Unexpectedly, the inhibition of STAT5 led to notable advantages in terms of cellular survival, with increase expression of Bcl2, an anti-apoptotic molecule and key indicator of cellular viability (**Fig 3I**). Together, our findings demonstrate that IL-2 and IL-15 favor the acquisition of exhaustion-like features at the expense of stem-like biology in *in vitro* conditions of repetitive TCR stimulation. In contrast, specifically targeting STAT5 activity with small molecule inhibitors restrains these unwanted cytokine effects and increase the quantity and quality of T-cell products destined for ACT, by promoting stem-like features.

**Fig. 3.**
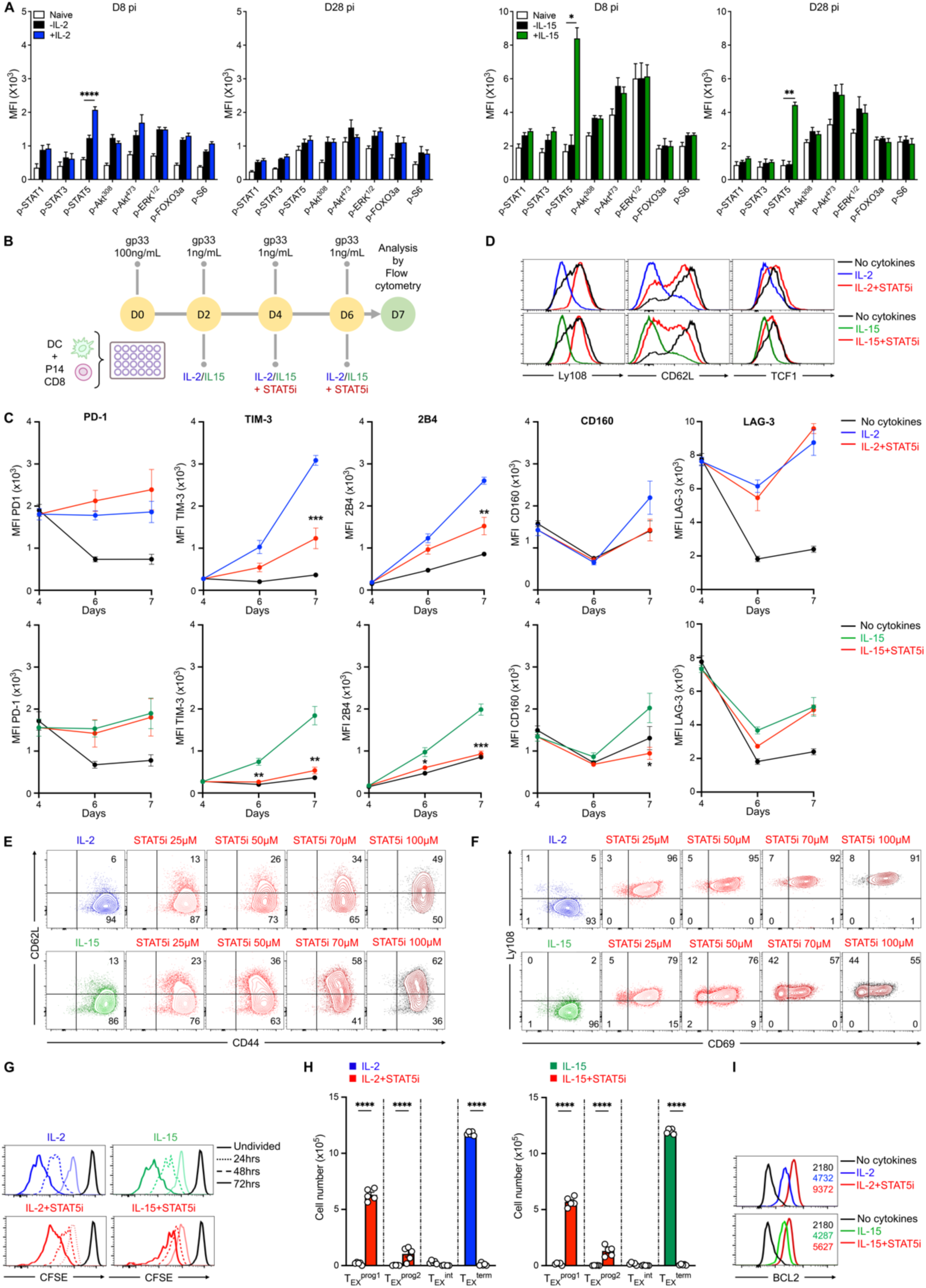
STAT5 inhibition favors the development of progenitor exhausted T cells *in vitro*. **(A)** Bar graphs of signaling proteins in CD8^+^ T cells from Cl.13-infected mice at day 8- and 28- pi following *ex vivo* IL-2 (left, blue) or IL-15 (right, green) stimulation (n=2 independent experiments, 24 mice). **(B)** Schematic representation of the experimental approach where P14 cells are co-cultured with DCs pulsed with gp33 alone (no cytokine), or combined with IL-2 (100 ng/mL; blue) or IL-15 (200 ng/mL; green) and where STAT5 inhibitor (STAT5i; red) is added to the culture on days 4 and 6 (100 μM unless otherwise specified). (**C**) MFI of indicated IRs at day 4, 6 and 7. (**D**) Histograms of Ly108, CD62L and TCF1 expression at day 7. **(E, F)** Contour plots of CD44/CD62L and CD69/Ly108 co-expression at day 7. **(G)** CFSE dilutions of day 4-expanded cells after 24, 48, and 72 hours. (**H**) Bar graphs of the absolute counts of T_EX_^prog1^ (CD69^+^Ly108^+^), T_EX_^prog2^ (CD69^-^Ly108^+^), T_EX_^int^ (CD69^-^Ly108^-^), and T_EX_^term^ (CD69^+^Ly108^-^) populations at day 7. (**I**) Histogram of BCL2 expression at day 7. Data are representative of 2-5 independent experiments (**D, E, F, G** and **I**) or combined from 3 independent experiments (**C** and **H**). Error bars indicate means ± SEM. *P < 0.05; **P < 0.01; ***P < 0.001; ****P < 0.0001, using Mann-Whitney test (**A**), two-way ANOVA (**C**) or one-way ANOVA (**H**).

### STAT5 inhibition supports a stem-like transcriptional program

To understand the underlying molecular impact of STAT5 inhibition on *in vitro* expanded CD8^+^ T cells, we performed RNAseq and compared P14 CD8^+^ T cells expanded *in vitro* with IL-2 or IL-15, with or without STAT5 inhibition, isolated on day 7 of culture. Naïve CD8^+^ T cells were also analyzed as a reference. A total of 211 and 351 genes were upregulated, and 71 and 246 genes were downregulated when comparing IL-2+STAT5i (IL2S5i) to IL-2 and IL-15+STAT5i (IL-15S5i) to IL-15, respectively (log2FC ≥ 0.5, and FDR < 0.05, **Fig 4A-B, Table S4-S5**). Notably, we observed concordance in the gene programs altered by STAT5 inhibition for both the IL-2 and IL-15 conditions (**Fig 4C**). This suggested that, at least in these *in vitro* settings, STAT5 triggers a similar program downstream of both IL-2 and IL-15. Differential gene expression analysis (DGEA) revealed that genes associated with T-cell memory and/or T-cell stemness, including *Sell* (CD62L), *Slamf6* (Ly108), *Ccr7*, *Tcf7* (TCF-1), *Id3*, and *Bach2*, *w*ere upregulated in the presence of STAT5i compared to both IL-2 or IL-15 conditions alone (*29*) (**Fig 4D**). Genes encoding critical proteins regulating egress, migration and/or homing of memory CD8^+^ T cells and resident memory T cells were also upregulated such as *S1pr1*, *Klf2* and *Itgae* (CD103). Genes associated with cytotoxic effector molecules such as Gzma, *Gzmb*, *Prf1* (Perforin 1), and Tnfsf10 (TRAIL) were downregulated in IL-2S5i- and/or IL-15S5i-expanded CD8^+^ T cells. Additionally, genes related to terminal exhaustion, including *Havcr2* (TIM-3) and *Entpd1* (CD39), were also downregulated in these cells (*30*).

**Fig. 4.**
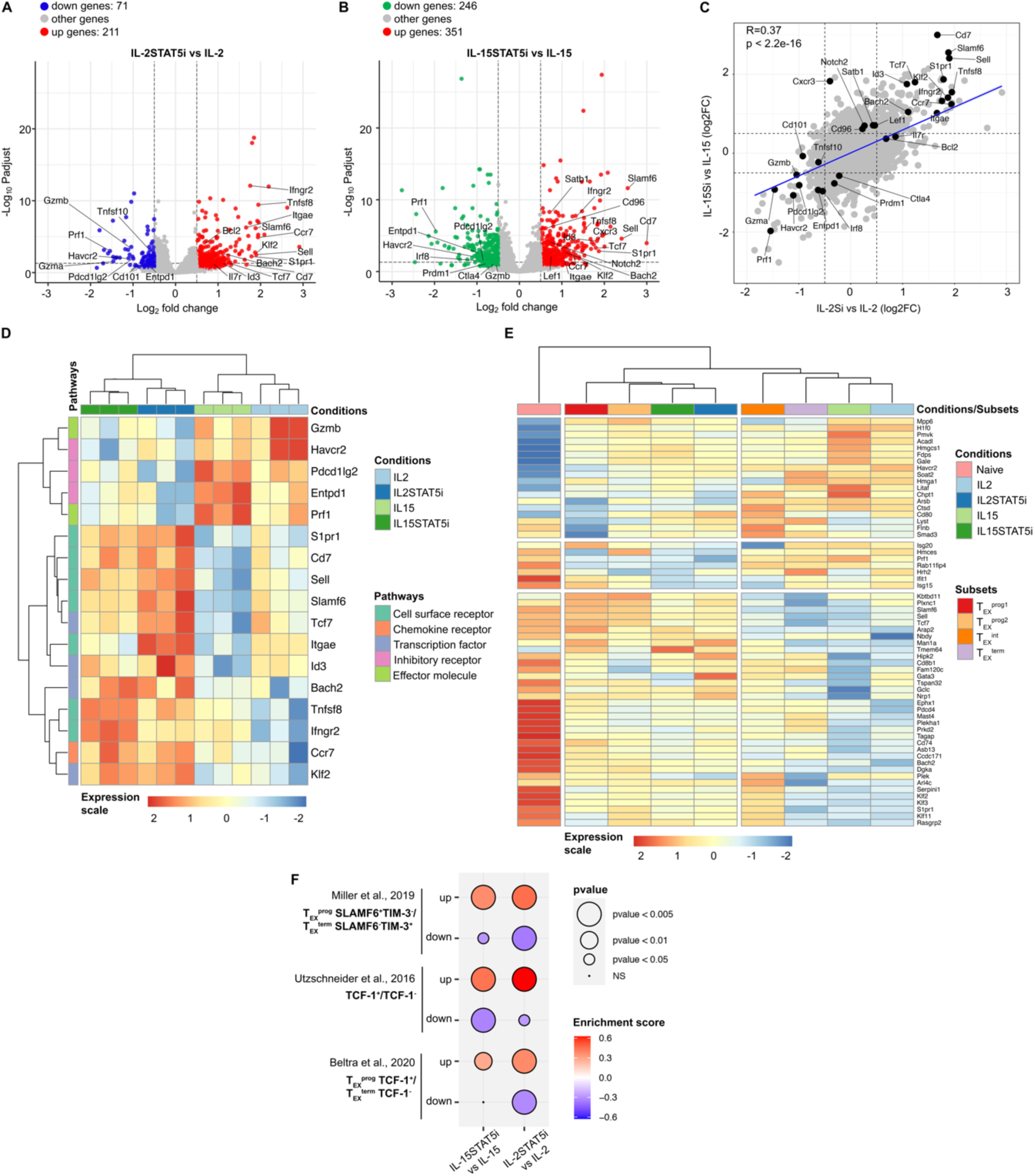
Molecular signature of CD8^+^ T cells stimulated with IL-2/IL-15 and STAT5 inhibitor reveals increased stemness following inhibition of STAT5. Volcano plots of downregulated genes (blue for IL-2 and green for IL-15) and upregulated (red) in (**A**) IL-2+STAT5i (IL-2S5i) versus IL-2 or (**B**) IL-15+STAT5i (IL-15S5i) versus IL-15 stimulated CD8^+^ T cells at day 7 of the culture (log₂ FC ≥ 0.5 and FDR < 5%, n=3 biological replicates). Labels indicate DE genes associated with T-cell exhaustion or stemness. **(C)** Scatterplot showing DE genes in IL-2+STAT5i vs IL-2 (X axis) and IL15+STAT5i vs IL-15 (Y axis) culture conditions. DE genes (black dots) associated with T-cell exhaustion or stemness are labelled. **(D)** Heatmap analysis displaying the hierarchical clustering of experimental conditions (n=3 per condition, X axis) alongside the top DE genes from **A-C** associated with T-cell exhaustion or stemness (right Y axis). Gene expression was log-normalized and expression-scaled across all samples, with genes categorized into pathways (left Y axis). **(E)** Heatmap analysis showing DE genes in naive and cytokine-stimulated CD8^+^ cells +/- STAT5i compared to genes from T_EX_^prog1^, T_EX_^prog2^, T_EX_^int^, T_EX_^term^ exhausted T cells from Beltra *et al.* (GSE149876) (*23*). **(F)** GSEA of DE gene profiles comparing IL-2- or IL-15-STAT5i vs IL-2- or IL-15-stimulated CD8^+^ T cells against T-cell progenitors versus T-cell exhausted gene signatures from published datasets (*19, 23, 31*) (p < 0.05, p < 0.01, p < 0.005).

The RNA-seq analysis also revealed that CD8^+^ T cells expanded with IL-2 and IL-15 were transcriptionally closer to the more differentiated exhausted subsets, specifically T_EX_^int^ and T_EX_^term^ populations, observed in chronically infected mice (**Fig 4E**, right side) (*23*). In contrast, CD8^+^ T cells treated with the STAT5 inhibitor displayed transcriptional profiles more similar to T_EX_^prog1^ and T_EX_^prog2^ progenitors (**Fig 4E**, left side). GSEA further showed positive enrichment of gene sets (red) associated with Slamf6^+^Tim-3^−^ progenitor-exhausted T cells (*19*), TCF1^+^ memory-like T cells (*31*), and TCF1^+^ progenitor T_EX_ cells (*23*) from chronic LCMV infection, in IL-2 or IL-15 plus STAT5i-treated cells (**Fig. 4F**). Conversely, genes associated with terminally exhausted Slamf6^−^Tim-3^+^ (*19*), TCF1^−^ T cells (*31*) and TCF1^-^ terminally exhausted T cells (*23*) were negatively enriched (blue) in STAT5i treated CD8^+^ T cells. Together, these observations confirm that suppression of STAT5 signaling downstream of IL-2 and IL-15 supports stem-like programs in *in vitro* expanded CD8^+^ T cells.

### Cytokine production and anti-tumor activity of CD8^+^ T cells is amplified by STAT5 inhibition

Repetitive Ag-signals during *in vitro* expansion protocols not only drive programs that resemble that of *in vivo* exhausted CD8^+^ T cells, but these cells also undergo substantial functional impairment (*32, 33*). The ability of STAT5 inhibition at reducing some features of exhaustion *in vitro*, as well as the demonstrated impact it had on the transcription of effector molecules such as granzymes, perforin and TRAIL, prompted us to evaluate functionality of CD8^+^ T cells expanded in settings of tempered STAT5 signaling. To do so, P14 CD8^+^ T cells expanded for a week under constant TCR-signaling and IL-2 or IL-15, in the presence or not of STAT5i, were re-stimulated for 6h at the end of the culture (day7) with a murine EL4 cell-line expressing the cognate LCMV D^b^gp33-41 peptide (EL4-gp33). We found that cells treated with STAT5i (under either IL-2 or IL-15 conditions) produced significantly more IFNγ compared to their non-treated counterpart (IL-2 or IL-15 alone) and expressed higher levels of the degranulation marker CD107 whereas the overall production of TNF-α and IL-2 appeared to be comparable across all conditions (**Fig. 5A-B**). More detailed analysis revealed that STAT5 inhibition impacted the quality of effector cells on a per-cell basis, with increased frequency of cells expressing high levels of TNFα, IFNγ and CD107a, and increased frequency of polyfunctional cells secreting all three effector proteins simultaneously (**Fig. 5C**). In contrast, and as expected from the transcriptomic data, STAT5i-treated cells, particularly in context of IL-15 stimulation, had reduced but not absent expression of granzyme B, in terms of both overall levels and frequency of high cells (**Fig. 5A-C**), suggesting potential impairment in immediate cytolytic potential.

**Fig. 5.**
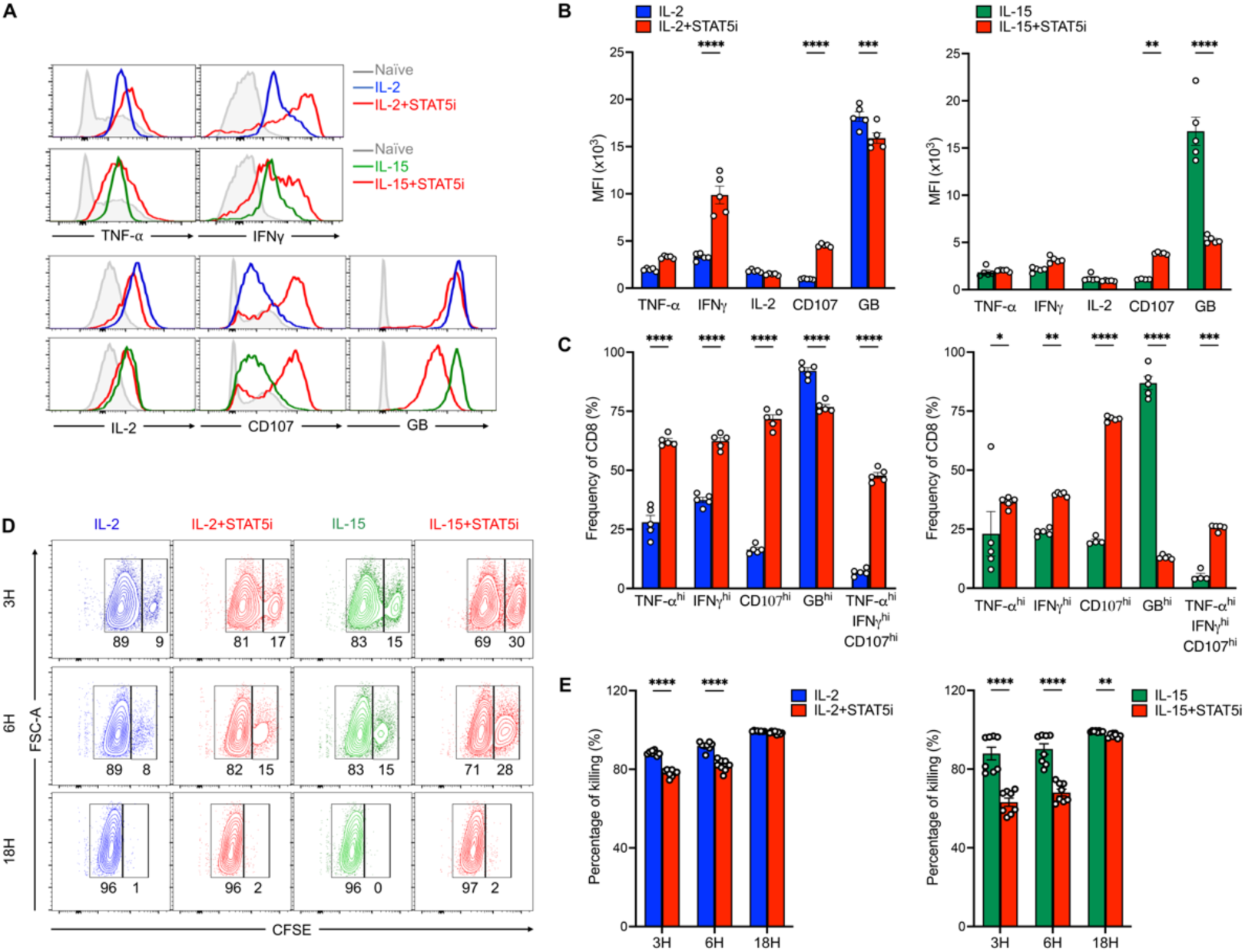
Inhibition of STAT5 enhances effector cytokine production and preserves cytotoxicity of T cells. P14 CD8^+^ T cells were expanded *in vitro* with continuous gp33 peptide and IL-2 (100 ng/mL; blue) or IL-15 (200 ng/mL; green) in the presence or absence of STAT5i (red), as previously described in Figure 3B, and harvested on day 7. Representative (**A**) histograms and (**B**) bar graphs showing the MFI of the indicated cytokines. (**C**) Bar graphs showing the frequency of cells expressing high levels of the indicated cytokines. **(D, E)** In vitro cytotoxicity assay where day 7 cells were co-cultured with CFSE-labelled EL4-gp33 (CFSE^high^, target cells) and EL4 (CFSE^low^, non-target cells) at a 1:1 effector-to-target ratio for 3, 6 or 18 hours. **(D)** Representative flow cytometry plot of the frequency of CFSE^high^ and CFSE^low^ cells at indicated time points. **(E)** Bar graphs showing the percentage of EL4-gp33 target cells lyzed by P14 CD8^+^ T cells expanded *in vitro* for 7 days. Data are representative of three independent experiments. Error bars indicate means ± SEM. *P < 0.05; **P < 0.01; ***P < 0.001; ****P < 0.0001, using one-way ANOVA (B and C) or two-way ANOVA (E).

To directly test cytotoxicity, we performed a time-course killing assay on P14 CD8^+^ T cells isolated after a week of *in vitro* expansion. Here, an equivalent number of P14 CD8^+^ T cells expanded in IL-2 or IL-15 (supplemented or not with STAT5i) were co-cultured (E/T ratio 1:1) with a 1-1-1 mixture of EL4-gp33 and EL4 cells not expressing the target antigen gp33 (EL4) previously labelled with high (CFSE^high^) or low (CFSE^low^) concentrations of Carboxyfluorescein succinimidyl ester (CFSE) respectively. Killing of CFSE^high^ EL4-gp33 was assessed at 3, 6 and 18h of co-culture. After 3 and 6 hours of co-culture, STAT5i-treated CD8^+^ T cells exhibited reduced killing of target cells compared to IL-2 or IL-15 conditions, consistent with the reduced expression of cytolytic molecule granzyme B in the former populations (**Fig 5D-E**). However, by 18h of co-culture, we observed near complete elimination of EL4-gp33 target cells across all conditions, indicating that all groups achieved similar levels of target cell elimination at that time point. This demonstrate that, despite a reduction in immediate killing, CD8^+^ T cells expanded in conditions of attenuated STAT5 signaling retain potent cytotoxic potential and exhibit effective target cell killing, while increasing their degranulation and cytokine secretion capacities.

We next sought to evaluate the anti-tumor potential of CD8^+^ T cells expanded in the context of limited STAT5 signals. We first tested their functional capacitites in a prophylactic setting by transferring OT1 CD8^+^ T cells (specific for the ovalbumin peptide OVA_257-264_) expanded for 7 days with the cognate peptide and IL-2, in the presence or not of STAT5i, into C57BL/6 mice one day prior to orthotopic injection of B16 melanoma cells expressing OVA_257-264_ (B16-OVA). (**Fig 6A**). Here, mice that received OT1 cells expanded with IL-2 or IL-2 + STAT5i exhibited delayed tumor growth compared to those that did not receive T cells (p values: 0.0068 and <0.0001 respectively) (**Fig. 6B**). ACT-treated mice also demonstrated prolonged survival compared to controls (median survival of 28 days versus 22 days) (**Fig. 6C**). Importantly, ACT with IL-2+STAT5i-treated cells more prominently reduced tumor growth than ACT with IL-2 (p: 0.079), and conferred the highest protection with increased survival of tumor-bearing mice (p: 0.0401) and 22% mice survival rate at day 60 (**Fig. 6 B-C**). All animals from the other groups, including mice receiving IL-2-expanded ACT, eventually succumbed to the tumor challenge. This suggested a protective advantage of STAT5i-expanded CD8^+^ T cells in controlling tumor growth, at least in the prophylactic setting.

**Fig. 6.**
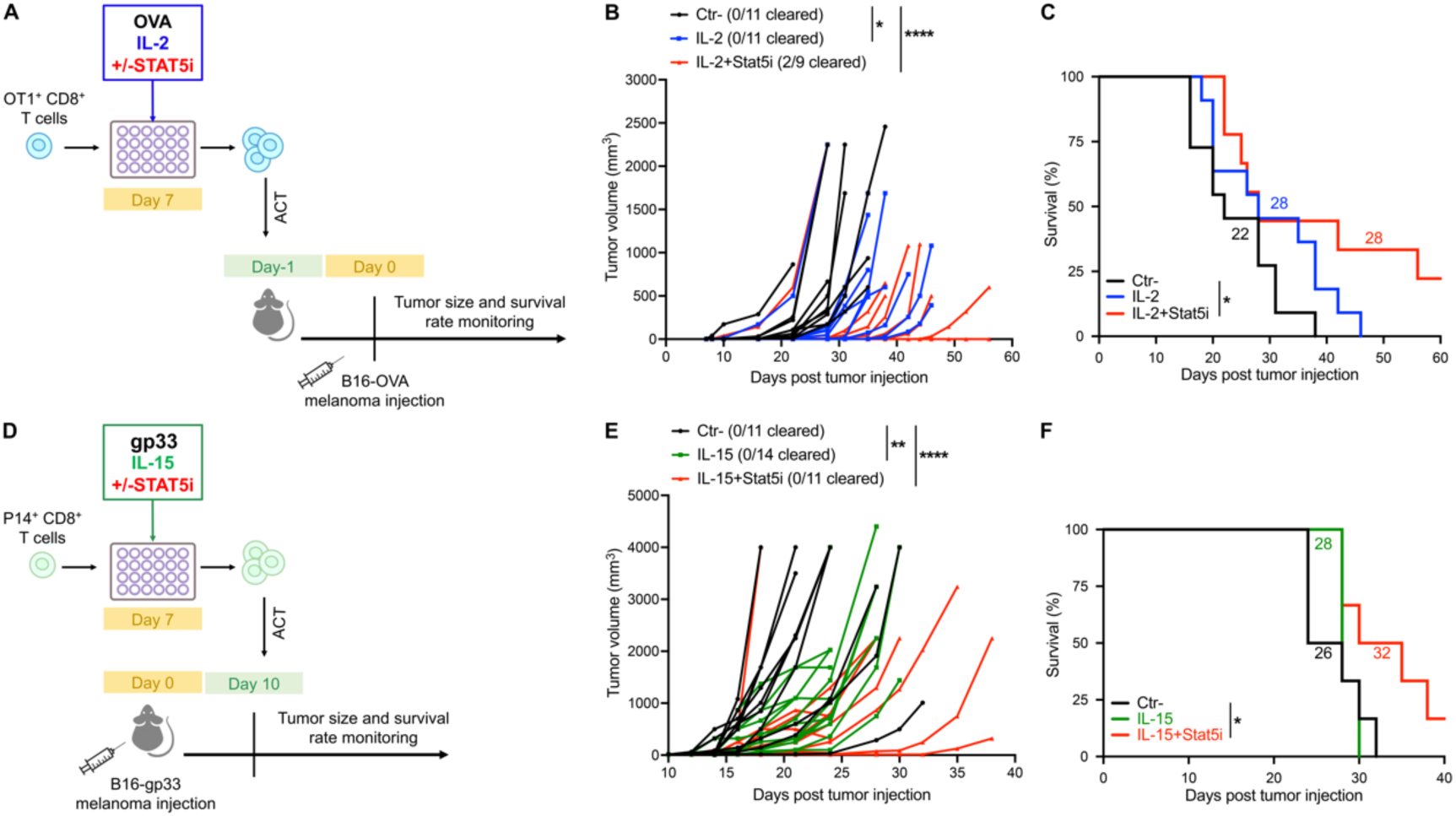
Adoptive transfer of CD8^+^ T cells expanded in the presence of STAT5i significantly reduces tumor size and enhances survival in mice. (**A**) Schematic of the experimental design. OT1^+^ CD8^+^ T cells stimulated with OVA, IL-2 (100ng/mL; blue line) +/- STAT5i (100μM; red line) for 7 days were adoptively transferred in B6 mice prior to their IV injection with 2 × 10^5^ B16-OVA tumor cells. Tumor size and animal survival was monitored. (**B**) Individual tumor growth and (**C**) mice survival following B16-OVA inoculation. (**D**) Schematic of the experimental design. B6 mice were inoculated with 2 × 10^5^ B16-gp33 tumor cells. P14^+^ CD8^+^ T cells stimulated *in vitro* for 7 days with continuous gp33, IL-15 (200ng/mL; green line) +/- STAT5i (100μM; red line) were adoptively transferred in tumor-bearing mice at day 10. Tumor size and animal survival was monitored. (**E**) Individual tumor growth and (**F**) mice survival curve after B16-gp33 cell injection. Data in (**B** and **C**) are combined from two independent experiments with 9-11 mice. Data in (**E**) is combined from two independent experiments with 11-14 mice. Data in (**F**) is representative of one of two experiments with 6-7 mice per group. Statistical significance was determined using two-way ANOVA (B and E) and the log-rank (Mantel-Cox) test (C and F). Median survival days are indicated on each graph.

We next evaluated the functional capacities of *in vitro* expanded CD8^+^ T cells in *bona fide* therapeutic settings when a tumor is already established. Here, P14 CD8^+^ T cells (specific for the glycoprotein (GP) peptide 33-41) were expanded for 7 days with its cognate peptide and IL-15, in the presence or absence of STAT5i and injected at day 10 post-tumor inoculation in B16-GP33 tumor bearing mice (**Fig. 6D**). Consistent with our prior results, CD8^+^ T cells expanded under attenuated STAT5 conditions had the greatest effect in reducing tumor growth (p <0.0001 for IL-15+STAT5i versus p:0.0047 for IL-15) (**Fig. 6E**). Similarly, mice that received IL-15 + STAT5i-treated cells exhibited prolonged survival (with a median of 32 days) compared to mice that received IL-15 treated cells (28 days) or controls (26 days) (p=0.0447 between STAT5i-treated cells and controls). Notably, only the IL-15 + STAT5i group had mice surviving beyond 40 days, with a survival rate of 16%, while all the other mice died by day 32 (**Fig. 6F**). Together, these experiments reveal that transient STAT5 inhibition during *in vitro* CD8^+^ T cell expansion enhances the *in vivo* efficacy and anti-tumor potency of adoptively transferred cells.

### STAT5 inhibition supports stemness and limits the exhaustion of human CAR T cells

Finally, we questioned if the beneficial impact of reduced STAT5 signaling on CD8^+^ T cell development towards increased stemness would be similarly beneficial in the human context. To investigate this question, we chose to manipulate STAT5 activity during the expansion of human CAR T-cells using protocols that favor CAR T-cell exhaustion (*34*). Here, CD8^+^ T cells isolated from human PBMCs were activated with aCD3/CD28 for 2 days and transduced with a CAR construct targeting the human antigen CD22 expressed by leukemic cells (*35–37*). CAR expressing cells were further expanded for an additional 12 days in complete media supplemented with aCD3/CD28 and IL-2 or IL-15. On day 7 and onward, STAT5i was added or not to the culture media every 2 days (**Fig. 7A**). Using CD39 and CD69 co-expression and the expression of IR to evaluate the phenotypic characteristics of the cells, we showed that human CD22-CAR T cells expanded in the presence of IL-2 and IL-15 differentiated into a CD39^+^CD69^+^ highly differentiated subset (**Fig. 7B-D**) and expressed high levels of PD1, Tim-2, 2B4 and CD39 (**Fig. 7E-F**). In contrast, CAR T cells expanded in the presence of IL-2/IL-15 and restricted STAT5 signaling displayed reduced expression of CD69, an early activation marker and CD39, an ecto-ATPase expressed by effector memory and exhausted T cells (**Fig. 7B**). This led to the generation of increased frequency and number of memory-progenitor stem-like cells (CD39^-^CD69^-^) (**Fig. 7C-D**), which persist and mediate anti-cancer response in melanoma patients treated with ACT (*4*). Other markers of memory cells where similarly increased in the presence of STAT5i, including the L-selectin CD62L (**Fig. 7E**), while the expression of IR was limited in this setting (**Fig. 7E-F**). Increased memory-progenitor differentiation and limited IR expression also correlated with restricted CAR T cell expansion after day 10 of the culture (**Fig. 7G**). To confirm that transient STAT5 inhibition did not restrict the functional anti-cancer potential of the expanded CAR T cells, we performed a typical cytotoxic assay. To do so, we co-cultured IL-2 expanded CD22 expressing CARs in the presence or absence of STAT5i with CD22 leukemic targets for 24 hours. Using various effector-to-target ratios, we demonstrated that CD22 CAR T cells expanded with or without STAT5i efficiently eliminated leukemic targets, particularly at high effector-to-target ratios (**Fig. 7H**). Together, these experiments indicate that transient STAT5 inhibition during human CAR T cell expansion favors memory-progenitor stem-like differentiation with preserving the anti-cancer cytotoxic potential of the CARs. Collectively, our results demonstrate that constant exposure to IL-2 and IL-15 cytokines skew the differentiation of murine and human CD8^+^ T cells towards an exhausted state, a state that can be reversed towards heightened stemness with enhanced anti-tumor functions by the transient inhibition of STAT5.

**Fig. 7.**
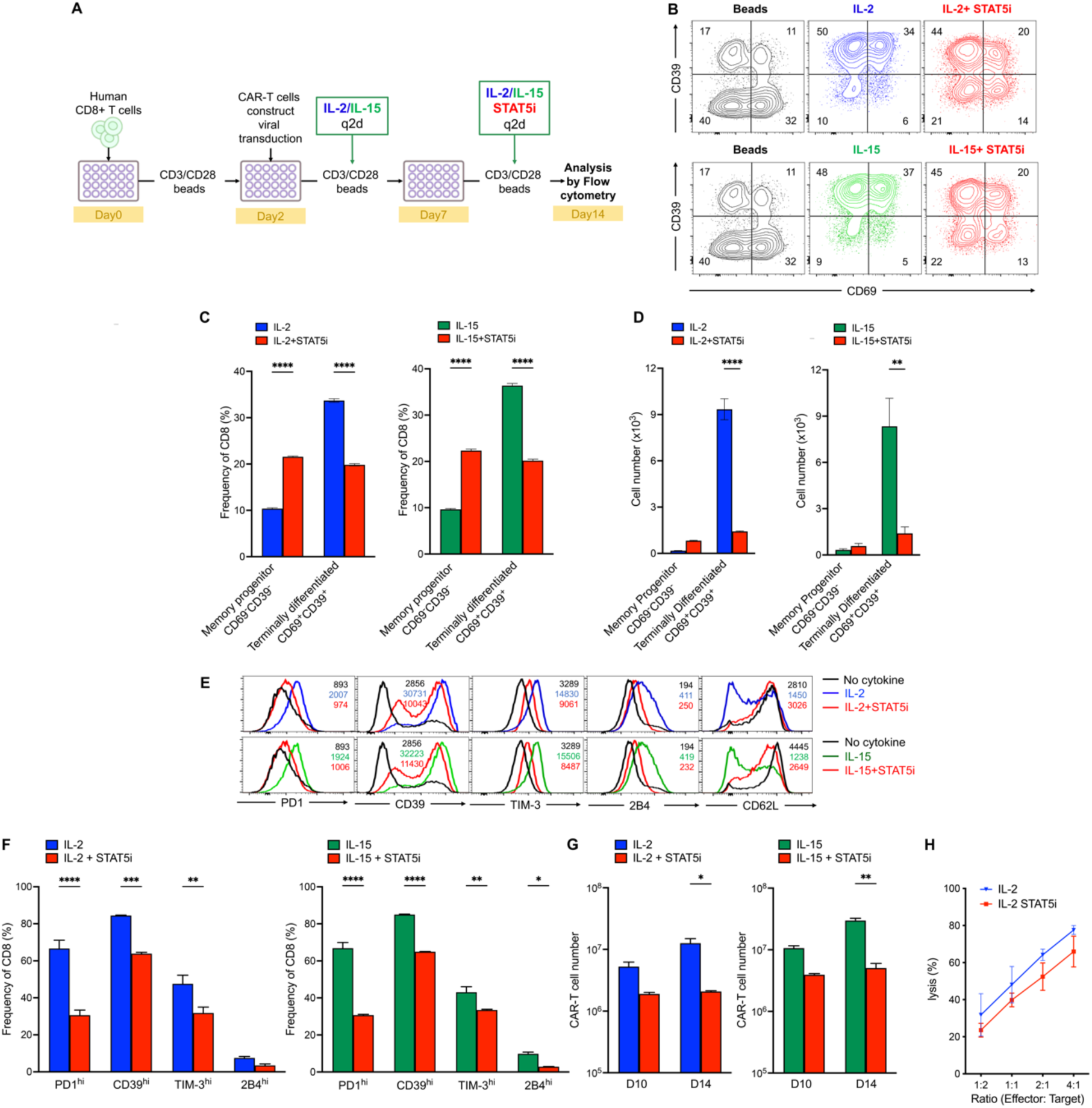
STAT5 inhibition favors the development of memory progenitor CAR-T cells with stem-like features. (**A**) Schematic of the experimental design. Human CD8^+^ T cells were enriched from PBMCs of healthy donors and activated using human CD3/CD28 Dynabeads for two days prior to transduction with an anti-CD22 CAR. CAR-T cells amplified for 2 weeks using CD3/CD28 beads and either human IL-2 (0.5 μg/mL; blue) or human IL-15 (0.5 μg/mL; green) every 2 days, +/- STAT5 inhibitor (100 μM; red) from day 7 onwards. (**B**) Representative flow cytometry plots of CD69/CD39 co-expression at day 14 of the culture. Bar graphs showing the frequency (**C**) and cell number (**D**) of memory progenitors (CD69^-^ CD39^-^) and terminally differentiated cells (CD69^+^ CD39^+^) cells. (**E**) Representative histograms of the indicated surface markers. (**F**) Bar graphs showing the frequency of cells expressing high levels of the indicated IRs. (**G**) Bar graphs showing the total CAR-T cell number obtained at day 10 and 14 of the culture when stimulated with IL-2 (blue) or IL-15 (red) in the presence (red) or not of STAT5i. (**H**) Percentage of EGFP^+^ CD22^+^ target cells lyzed in 24 hours by CAR-T cells stimulated with IL-2 +/- STAT5i at indicated effector:target ratios. Data in (B-G) are representative of two independent experiments. Data in (H) are combined from 3 independent experiments with nine technical repeats. Error bars indicate means ± SEM. *P < 0.05; **P < 0.01; ***P < 0.001; ****P < 0.0001, Using 2Way ANOVA.

## DISCUSSION

To enhance the efficacy of ACT, it is essential to thoroughly understand the role of cytokine signaling in T-cell exhaustion. We implemented an *in vitro* protocol to expand T cells, enabling us to investigate how cytokines influence their differentiation and exhaustion. Our results indicate that inhibiting IL-2/IL-15 signaling via STAT5 promotes the development of stem-like progenitors and prevents terminal exhaustion in mouse CD8^+^ T cells and human CAR T-cell expansion protocols *in vitro*. Additionally, STAT5 inhibition enhances tumor control in mice following the injection of STAT5i-treated CD8^+^ T cells *in vivo*.

Cytokines have been shown to regulate T-cell responses and are critical modulators in conditions of chronic TCR stimulation, as in chronic viral infection, cancer and *ex vivo* cell therapy expansion (*8, 38*). IL-6, IL-10, TNF-α and TGF-β are well known contributors to T-cell exhaustion, and blocking their actions improves anti-viral and anti-cancer immunity (*39–41*). Cytokines of the γ_c_ family are also crucial regulators of CD8 T-cell survival, proliferation and differentiation (*42*). In chronic viral infections, we have shown that IL-2 and IL-15 through IL2Rβ support the expression of key inhibitory receptors (*18*). Here, we demonstrate that blocking IL2Rβ signaling *in vivo* supports stemness over exhaustion at both the transcriptional and protein levels. In ACT protocols, IL-2 is typically used to expand T cells in sufficient numbers (*43*). Unfortunately, this approach promotes the accumulation of terminally differentiated cells, as demonstrated here by the T-cell exhaustion signature of cells expanded with IL-2 (*27, 28, 44*). Interestingly, IL-15, which shares the IL2Rβ chain with IL-2, has been proposed for use in ACT protocols. Unlike IL-2, IL-15 may support T-cell expansion while preserving cell functionality. Given the close similarities between these two cytokines - signaling through near identical quaternary structures and kinases, and comparable strength-dependent signaling kinetics, we questioned if IL-15 also promoted T-cell exhaustion in context of chronic TCR stimulation. Here, we showed that high doses of IL-15, similar to IL-2, promoted the expression of inhibitory receptors and the terminal differentiation of exhausted T cells (T_EX_^term^). This is reminiscent of what has been observed in human NK cells exposed to prolonged IL-15 stimulation (*45*) and contrasts with IL-15’s capacity to promote memory homeostasis when used at low concentrations (*46*). Indeed, using a stringent *in vitro* model with specific concentrations of peptide and cytokine, we were able to show that two closely-related cytokines, i.e. IL-2 and IL-15, impart similar transcriptional programs in T-cells when used in comparable and progressively increased concentrations. Importantly, this *in vitro* model allowed us to interrogate the signaling molecules and transcription factors that regulate T-cell exhaustion over stemness in a controlled fashion. Such *in vitro* protocol will be particularly useful in the future to perform *in vitro* screens to reduce functional exhaustion and/or validate therapeutic targets that support stemness. Notably, a comparable *in vitro* model was recently reported through transcriptional and epigenetic studies as a great model to interrogate mechanisms driving T_EX_ differentiation (*32*).

Using our controlled *in vitro* protocol, we investigated the signals downstream of IL2Rβ, which we have shown to be a critical regulator of T-cell exhaustion in both humans and mice (*18*). IL-2 and IL-15 through IL2Rβ may signal through multiple pathways. Our chronic viral infection model suggested that pSTAT5, rather than other signaling pathways such as RAS-MAPK, PI3K-mTORC1, and PI3K-AKT, was particularly critical in context of chronic antigenic stimulation. Thus, we employed our *in vitro* protocol to depict the signaling cascade that regulate T-cell exhaustion in response to IL-2 and IL-15. We showed that the inhibition of STAT5 or JAK3 with small molecule inhibitors during the expansion of murine CD8^+^ T cells or human CAR-T cells led to increased number of cells with characteristics of stem-like progenitor cells and increased anti-tumor functions. This supports previous report highlighting the crucial role of STAT5 inhibition in reducing IRs expression on CD8^+^ T cells (*27*). We have showed that the phenotypic changes observed were dose- and inhibitor-dependent, reinforcing the importance of inhibitor selection and concentration in ACT protocols. Interestingly, a recent study has demonstrated that STAT5 plays a pivotal role in sustaining effector CD8^+^ T cell responses during chronic viral infection. Specifically, STAT5 is crucial for enabling the transition from Tex^prog^ to Tex^int^ cells (*47*). Our results demonstrate that increased stemness is favored in contexts of reduced STAT5 signaling and that transient inhibition of STAT5 does not hamper ultimate effector functions. In contrast, persistent and high dose stimulation with IL-2 and IL-15 may negatively hamper core T-cell functions as it will favor end-stage exhaustion. Altogether, this illustrates that effective CD8^+^ T cell differentiation depends on properly timed JAK-STAT signaling. Targeted disruption of IL-2Rβ signaling at precise stages of differentiation could provide valuable insights for enhancing the efficacy of ACT through increased stemness.

The stemness of T cells is crucial for ACT as it supports self-renewal, multipotency, proliferative capacity, and longevity (*48*). As a result, cell products with increased stemness potential provide prolonged immune surveillance and durable and potent immune responses against cancer and virally-infected cells (*49*). The concept of stemness extends from embryonic stem cells to cancer stem cells, with specific transcription factors varying according to stem cell type. STAT3, TCF1, LEF1, ID3, and BCL-6 were shown to be important promoters of the stemness of CD8+ T cells (*49, 50*). Here we show that inhibition of STAT5 or JAK3 support the development of stem cells through increased transcription of TCF1, ID3, BACH2 and KLF2. Interestingly, targeting other signaling cascades downstream of the TCR and/or cytokine receptors, such as the RAS-MAPK and PI3K-AKT pathways, were also reported to result in less differentiated T cells and improved antitumor activity (*51–53*). While it is unclear whether these results were solely due to increased stem cell numbers, both our findings and others strongly suggest that timely inhibition of signaling kinases may enhance the quality and potency of ACT products.

In conclusion, our study highlights the importance of IL-2 and IL-15 signals in driving T-cell exhaustion and underscores the importance of inhibiting STAT5 in a timely manner to enhance T-cell stemness. It also points to the complexity of targeting signaling molecules within a narrow therapeutic window to optimize the potency of ACT products while maintaining adequate T-cell numbers and functionality. Despite these challenges, our findings suggest that transient STAT5 inhibition during ACT expansion could be a promising strategy to foster enhanced stemness and potent anti-cancer functions. Ultimately, our research underscores the critical importance of precisely adjusting cytokine doses and signaling pathways at key stages of T-cell development, paving the way for more effective and targeted cancer immunotherapies that can overcome current therapeutic limitations and greatly improve patient outcomes.

## MATERIALS AND METHODS

### Mice, viral infection and adoptive co-transfer

CD45.2^+^ P14^+^ TCR transgenic mice, which express a TCR specific for the LCMV H-2Dbbgp33-41 peptide were bred in-house and backcrossed onto the C57BL/6 (B6) (RRID: IMSR_JAX:000664) background, as previously detailed (*54*). For the adoptive transfers using the LCMV model, CD45.1^+^ B6 mice were purchased from Jackson Laboratory and bred with P14^+^ TCR transgenic mice to obtain CD45.1^+^ CD45.2^+^ P14^+^ IL-2Rβ^+/+^ mice. RAG2^-/-^ IL2Rβ-deficient P14^+^ TCR transgenic mice (IL-2Rβ^-/-^, CD45.2^+^) were obtained by breeding the above P14^+^ mice with RAG2^-/-^ mice and IL2Rβ^-/-^ mice from Jackson Laboratory as previously described (*18, 54*). All mice were housed in specific pathogen-free facilities at Sainte-Justine University Hospital (CHUSJ) Research Center in Montreal, Canada. Experiments were conducted under Canadian Council on Animal Care guidelines and were approved by the Institutional Committee for Animal Care at CHUSJ Research Center. LCMV Cl.13 was produced in BHK-21 cells (RRID: CVCL_1914) and titrated through plaque assays on Vero cells (RRID: CVCL_0059) as previously described (*18, 54*). To establish a chronic LCMV infection model, 2 × 10^6^ plaque-forming units (PFU) of LCMV CL.13 were injected intravenously (i.v.) into B6 mice. For adoptive co-transfer studies, 5,000 P14^+^ IL-2Rβ^+/+^ (CD45.1^+^ CD45.2^+^) and 5,000 P14^+^ IL-2Rβ^-/-^ (CD45.2^+^) cells were adoptively co-transferred into sex- and age-matched recipient B6 CD45.1 mice 24 h prior to Cl.13 infection. The phenotype of transferred P14^+^ cells was analyzed by flow cytometry 28 days p.i..

### Flow cytometry

Cell suspensions from spleens were stained with LIVE/DEAD Aqua (ThermoFisher Scientific, MA, USA) and antibodies against cell surface antigens for 30 minutes at 4 °C in phosphate buffered saline 1X (PBS 1X). Subsequently, cells were fixed and permeabilized at 4°C using Cytofix/Cytoperm (BD Biosciences, NJ, USA) for 30 minutes, followed by staining with intracellular antibodies in Perm/Wash buffer (BD Biosciences, NJ, USA) for 1 hour at 4°C. For *ex vivo* cytokine stimulation, cells were rested in complete RPMI for 4 hours and stimulated with 100 U/mL recombinant murine IL-2 or IL-15 (PeproTech, NJ, USA) in complete Roswell Park Memorial Institute (RPMI) 1640 supplemented with 10% heat-inactivated FBS, 2 mmol/L L-glutamine, 1 mmol/L sodium pyruvate, 1% nonessential amino acids, 25 mM/L HEPES, 100 U/mL penicillin-streptomycin, and 50 μM 2-mercaptoethanol) for 45 minutes prior to flow cytometry staining and analysis. For p-STAT staining, cells were fixed with 2% paraformaldehyde in PBS 1X for 30 minutes at room temperature, then fixed with ice-cold methanol for 30 minutes at 4 °C and stained for p-STAT antibodies for 1 hour in PBS 1X with 2% fetal bovine serum (FBS). H-2Db/gp33-41 biotinylated monomers were purchased from the NIH Tetramer Core Facility and coupled in-house to streptavidin R-phycoerythrin. Anti-mouse antibodies CD16/CD32 (2.4G2), CD8 (53-6.3), PD1(RMP1-30), CD160 (CNX46-3), CD3 (H57-597), LAG-3 (C9B7W), CD44 (IM7), TNF-α (MAb11), IFN-γ (XMG1.2), p-STAT1 (pY701), p-STAT3 (py705), and p-STAT5 (py694s) were obtained from BD Biosciences (NJ, USA). TIM-3 (RMT3-23), CX3CR1 (SA011F11), Ly108 (330-AJ), 2B4 (M2B4), IL-2 (4A9) and CD107 (1D4B) were sourced from Biolegend (CA, USA). CD69 (H1.2F3), TOX (TXRX10), CD101 (Moushi10), 2B4 (eBio244F4), CD45.1 (A20), CD45.2 (104), Granzyme B (NGZB) were provided from Invitrogen (CA, USA) and TCF1 (C63D9), p-AKT308 (D25E6), p-AKT473 (D9E), p-ERK(T202), p-S6 (D57.2.2E), and p-FOXO3 (S253) were supplied from Cell signaling (MA, USA). Data were acquired with an LSR FORTESSA II with High Throughput Sampler 192 (HTS) from BD Biosciences (MA, USA). FlowJo software 10.7.1 (RRID:SCR_008520) was used to perform all data analysis. Fluorescence Minus One (FMO) and unstimulated conditions were used to set the gates.

### *In vitro* CD8^+^ T-cell expansion

Mouse P14^+^ CD8^+^ T cells were expanded *in vitro*, as described previously (*18*). Briefly, 2 × 10^5^ P14^+^ CD8^+^ T cells were co-cultured with dendritic cells (1:1) isolated from B6 mice using EasySep^TM^ Mouse CD11c Positive Selection Kit II (Stemcell, MA, USA) with a high dose of gp33 peptide (100 ng/ml) in complete RPMI. At days 2, 4, and 6, P14^+^ CD8^+^ T cells were numbered, normalized, and re-cultured in complete RPMI with a lower dose of gp33 peptide (1 ng/ml) either alone or in combination with IL-2 (100ng/ml) or IL-15 (200ng/ml). To investigate the impact of JAK/STAT inhibitors, tofacitinib (Tofa, Selleckchem, TX, USA; 1μM), JAK3 inhibitor (Jacki, MilliporeSigma, MA, USA; 3 μM), or STAT5 inhibitor (STAT5i, MilliporeSigma, MA, USA; 100 μM) were added to the culture medium on days 4 and 6. On day 7, cells were collected, counted, and their phenotypes assessed by flow cytometry.

### *In vitro* killing, proliferation and cytokine production studies

For the *in vitro* killing assay, cells were harvested at day 7 of the culture, washed and then co-cultured with EL4-gp33 target cells (stained with 10 μM CFSE) and EL4 non-target cells (stained with 1 μM CFSE) at a 1:1 effector to target/non-target ratio for 3, 6, and 18 hours in complete RPMI medium. Specific killing was quantified using LIVE/DEAD Aqua staining and calculated as follows: 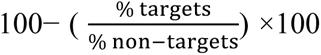. For intracellular cytokine staining, the *in vitro*-expanded P14 CD8^+^ T cells were collected on day 7 and re-stimulated with EL4-gp33 for 6 hours in the presence of Golgi Plug (BD Biosciences, NJ, USA). Cells were washed, stained as previously described for flow cytometry. In the proliferation assay, CD8^+^ T cells in the presence or absence of STAT5i were stained with carboxyfluorescein succinimidyl ester (CFSE, ThermoFisher Scientific, MA, USA) (10 μM) on day 4 of expansion. Their proliferation was monitored over the following 24, 48, and 72 hours by measuring CFSE dilution via flow cytometry.

### Adoptive T-cell transfer in melanoma mouse models

B16.F10 melanoma cells (BCRJ Cat# 0046, RRID: CVCL_0159) expressing the gp33 (B16gp33) or OT1 (B16Ova) peptides were cultured in Dulbecco’s Modified Eagle Medium (DMEM) supplemented with 10% FBS, 2 mM L-glutamine, and 100 U/mL penicillin-streptomycin, and maintained under G418 selection at 0.6 mg/mL (all reagents sourced from Life Technologies, CA, USA). Between 0.5 and 1 × 10^6^ P14^+^ CD8^+^ T cells, stimulated with IL-2 or IL-15 and cultured with or without STAT5i, were harvested on day 7 of culture and subsequently intravenously injected (i.v.) into B6 mice. Tumor cells (2 × 10^5^ cells in 200 mL PBS 1X) were subcutaneously injected one day prior (B16-Ova model) or 10 days after (B16-gp33 model) T-cell transfer. The survival was monitored daily, and the size of visible tumors was measured every three days using calipers to determine the longest and shortest diameters. Tumor volume was calculated using the formula: (length × width^2) / 2.

### Sample and library preparation for RNA sequencing (RNA-seq)

For the analysis of cells expressing high or low levels of the IL2Rβ chain following infection, total RNA was extracted from CD8^+^ Tet^+^ CD11a^+^ IL2Rβ ^hi^ or IL2Rβ ^lo^ T cells at day 21 post-LCMV Cl.13 infection, using the Weis lab RNA extraction protocol (*55*). Briefly, cells were sorted on a FACSAria flow cytometer cell sorter and directly isolated and lysed into Trizol for 5 min at room temperature. Phase separation was achieved by adding chloroform (1:4), followed by centrifugation at 12,000 x g for 15 minutes at 4°C. The aqueous phase containing RNA was transferred to a new tube, precipitated with isopropanol, and centrifuged at 12,000 x g for 10 minutes at 4°C to pellet the RNA. The RNA pellet was washed with 75% ethanol, centrifuged at 7,500 x g for 5 minutes at 4°C, air-dried, and resuspended in RNase-free water. Library preparation was carried out using the TruSeq Stranded mRNA kit according to the manufacturer’s instructions (Illumina, CA, USA). Poly-RNA sequencing was performed on the HISeq 2500 High Output V4 system. For the analysis of cells expanded in vitro in the presence of cytokines with or without STAT5i, RNA was isolated from naïve or expanded CD8^+^ T cells using miRNeasy Mini Kit (QIAGEN, Venlo, The Netherlands) following the manufacturer’s instructions. RNA samples were sent to the Institute for Research in Immunology and Cancer (IRIC) genomics platform in Montreal, Canada, for quality assessment and RNA sequencing. Poly-A mRNA enrichment was conducted using Dynabeads Oligo(dT) (ThermoFisher, MA, USA), and library preparation was carried out with the KAPA RNA Hyperprep Kit (Roche, Basel, Switzerland) using manufacturer’s instructions. RNA sequencing was performed using a NextSeq HighOutput 150 cycles kit (PE75) on the Nextseq500 (Illumina, CA, USA).

### RNA-sequencing analysis

All RNA-seq FASTQ files were aligned and mapped to the mouse genome (GENCODE M6/Ensembl v81 GRCm38 for the *in vivo* IL2Rβ^hi/lo^ experiment; GENCODE 31/Ensembl v97 GRCm38/mm10 for the *in vitro* IL-2/IL-15 with or without STAT5i experiments) using the STAR aligner (v2.4.2 for IL2Rβ and v2.7 for IL-2/IL-15; DOI: 10.1093/bioinformatics/bts635) with 2-pass mapping mode and a re-generated genome index. For the IL2Rβ^hi/lo^ analysis, reads were trimmed using *Cutadapt* (DOI: 10.14806/ej.17.1.200) to remove adapter sequences (GATCGGAAGAGCACACGTCTGAACTCCAGTCAC) and low-quality read ends (quality cutoff: 20) before alignment. The raw gene count matrices were generated using HTSeq (v0.6.1p1 for IL2Rβ^hi/lo^ and v0.13.5 for IL-2/IL-15 +/- STAT5i; DOI: 10.1093/bioinformatics/btu638) in non-stranded mode. Raw counts were filtered to remove low-expressed transcripts, retaining only protein-coding genes. For IL2Rβ^hi/lo^, additional filtering excluded RIKEN genes, prioritizing genes with clear one-to-one orthologous relationships to human genes. Differential expression analysis was conducted using DESeq, RRID:SCR_000154 (v1.26) in both cases, with its internal filtering step to detect and exclude outliers. Significance thresholds were defined as log2 fold change (FC) ≥ 0.5 and false discovery rate (FDR) < 5%, corrected by the Benjamini-Hochberg (BH) procedure. Raw gene count matrices from the public datasets GSE149876 (41), GSE83978 (38), and GSE123235 (52), which profile different subsets of progenitor and terminally exhausted CD8⁺ T cells during chronic LCMV Cl.13 infection, were downloaded from the Gene Expression Omnibus (GEO) (RRID:SCR_005012) and processed using the same pipeline as described for the IL-2/IL-15 with or without STAT5i experiments. For GSE149876 (Beltra *et al.*, day 30 post-infection) (*23*), subsets included T_EX_^prog1^ (Ly108⁺CD69⁺), T_EX_^prog2^ (Ly108⁺CD69⁻), T_EX_^int^ (Ly108⁻CD69⁻), and T_EX_^term^ (Ly108⁻CD69⁺). In GSE83978 (Utzschneider *et al.*, day 28 post-infection) (*31*), subsets were defined as exhausted phenotype (Tcf1^-^) and non-exhausted phenotype (Tcf1^+^). For GSE123235 (Miller *et al.*, day 30 post-infection) (*19*), progenitor exhausted cells were identified as Slamf6⁺Tim-3⁻ and terminally exhausted cells as Slamf6⁻Tim-3⁺. In all the Gene Set Enrichment Analysis (GSEA), the top 250 up- and down-regulated genes from the T-cell exhaustion signatures were mapped to the IL2Rβ^hi/lo^ and IL-2/IL-15 transcriptomic profiles. For **Figure 2F**, the GSEA analysis uses a T cell exhaustion gene signature from Bengsch *et al.*’s reanalysis of ATAC-Seq data (GSE86881(*56*), GSE97646)(*57*)). The upregulated and downregulated genes were obtained from *Bengsch et al.*’s supplementary table (NIHMS969093-supplement-1.xls), representing an exhaustion-specific gene list (murine and human orthologs)(*24*). All analyses were performed using custom R scripts along with additional packages (ggplot2 (RRID:SCR_014601), pheatmap, ggpubr, fsgea). For **Fig. 2E** and **Fig. 4D**, genes associated with T-cell exhaustion were categorized into pathways based on classifications from the Universal Protein Resource (UniProt).

### Human CAR-T cell culture and cytotoxicity assay

Human PBMCs were isolated from fresh blood of healthy donors using Ficoll-Paque density gradient centrifugation. CD3^+^CD8^+^ T cells were purified through negative magnetic selection using a Human CD8^+^ T Cell Enrichment Kit (Miltenyi Biotec, Bergisch Gladbach, Germany). Purified CD8^+^ T cells were stimulated with anti-Human T-Activator CD3/CD28 Dynabeads at a bead-to-cell ratio of 1:1 (ThermoFisher Scientific, MA, USA) in complete RPMI. Two days later, the cells were transduced with anti-CD22 CAR 28z that includes the 4-1BBz and CD3ζ signaling domains (m971 Myc tag 28z), as previously described (*36, 37*), and CAR expression was evaluated after 72 h using Flow cytometry. Subsequently, T cells were maintained in fresh medium, refreshed every two days and adjusted to a concentration of 1 × 10^6^ cells/mL. New stimulation were applied using new beads at a ratio 1:1 in combination with recombinant human IL-2 (0.5 μg/mL) or IL-15 (0.5 μg/mL, Peprotech, NJ, USA) for 14 days. Starting from day 7, STAT5i (100uM) was added to the culture medium every two days until day 14. Cells were collected on day 7 and 14 of expansion for flow cytometry staining and analysis as described above. For cytotoxicity assays, expanded CAR-T cells were co-cultured with EGFP pre-B-ALL cell line 697 CD22^+^ target cells at various effector to target ratios (4:1, 2:1, 1:1, 1:2) in RPMI supplemented with 2% FBS; 100 U/mL penicillin-streptomycin and 20 U/mL IL-2, for 24 hours. The number of surviving target cells was determined by flow cytometry. The percentage of lysis was calculated as follow: 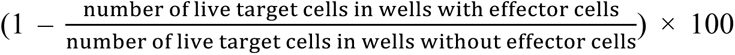. Ethical protocols for using human PBMCs were approved by the Research Ethics Board (REB) at the CHUSJ Research Center under study MP-21-2011-312. Written informed consent was obtained from all healthy donors.

### Statistical analysis

Statistical analyses were conducted with GraphPad Prism, RRID:SCR_002798. Statistical analysis of data was performed using Mann-Whitney and unpaired student’s t-test. Simultaneous comparisons of more than two groups were performed using one-way analysis of variance (ANOVA) or two-way ANOVA, as indicated in the figure legends. Statistical significance of survival curves was assessed using the log-rank (Mantel-Cox) test. Significance was set as *p <.05; **p <.01; ***p <.001; ****p <.0001. In all cases, P < 0.05 was considered statistically significant. Data in bar graphs and scatter plots are displayed as mean ± SEM.

## Acknowledgments

We thank Antonio Freitas for providing CD45.2^+^ P14 transgenic mice (Institut Pasteur, Paris, France), Rolf M. Zinkernagel for the initial LCMV Cl.13 strain (Zurich University Hospital, Zurich, Switzerland), Alain Lamarre (INRS-Institut Armand Frappier, Laval, Canada) and Martin Richer (McGill University, Montreal, Canada) for providing additional LCMV Cl.13 aliquots; and the NIH Tetramer Core Facility for the H-2Db/gp33-41 biotinylated monomers (Emory University Vaccine Center, Atlanta, USA). We also thank Philippe Bousso (Institut Pasteur, Paris) for the EL4-CFP cell line, Alain Lamarre (INRS-Institut Armand Frappier, Laval, Canada) for the B16.gp33 melanoma cells and Nathalie Labrecque (Institut de recherches cliniques de Montréal (IRCM), Montreal, Canada) for the B16.OVA cells. We are thankful to Luis Barreiro (University of Chicago, Illinois, USA) for help in the transcriptomic studies and critical review of the manuscript and Gabrielle Sutton for her technical assistance. Special thanks to all members of the animal facilities at CHUSJ Research Center, as well as Ines Boufaied from the flow cytometry platform, and Séverine Landais and Olivier Gingras from the Plateforme intégrée de génomique unicellulaire for their expertise and guidance.

This project was supported by the Canadian Institutes of Health Research (CIHR) awarded to HD (*Funding Reference umber: PJT-175209*). HD is also a Fonds de Recherche du Québec – Santé (FRQS) Clinical Research Scholar. MS and CR were supported by Cole Foundation Postdoctoral and Doctoral Scholarships respectively. J-C.B is supported by grants from the Swiss National Science Foundation (TMSGI3-218028) and the Parker Institute for Cancer Immunotherapy (PICI).

## Author contributions

Design of the experiments: MS, HD

Performed the experiments: MS, CR, AV, HR, EK

Biological data analyze: MS, CR, AV, BB

Bioinformatic analysis: NH

Supervision: HD, VPL

Figure formatting: BB, MS

Writing – original draft: MS, BB

Writing – review & editing: HD, JCB, EH, VPL

## Conflict of interest

The authors declare no competing interests.

## Data and materials availability

Data are available upon request. Data analyses from the RNAseq are available in Supplementary Tables S1 to S5.

## Notes

### Competing Interest Statement

The authors have declared no competing interest.

## REFERENCES

1. S. Guedan, M. Ruella, C. H. June, Emerging Cellular Therapies for Cancer. Annu Rev Immunol 37, 145–171 (2019).

2. R. C. Sterner, R. M. Sterner, CAR-T cell therapy: current limitations and potential strategies. Blood Cancer J 11, 69 (2021).

3. R. J. Kishton, S. K. Vodnala, R. Vizcardo, N. P. Restifo, Next generation immunotherapy: enhancing stemness of polyclonal T cells to improve anti-tumor activity. Curr Opin Immunol 74, 39–45 (2022).

4. S. Krishna, F. J. Lowery, A. R. Copeland, E. Bahadiroglu, R. Mukherjee, L. Jia, J. T. Anibal, A. Sachs, S. O. Adebola, D. Gurusamy, Z. Yu, V. Hill, J. J. Gartner, Y. F. Li, M. Parkhurst, B. Paria, P. Kvistborg, M. C. Kelly, S. L. Goff, G. Altan-Bonnet, P. F. Robbins, S. A. Rosenberg, Stem-like CD8 T cells mediate response of adoptive cell immunotherapy against human cancer. Science 370, 1328–1334 (2020).

5. V. Janelle, J. S. Delisle, T-Cell Dysfunction as a Limitation of Adoptive Immunotherapy: Current Concepts and Mitigation Strategies. Cancers (Basel) 13, (2021).

6. L. Gattinoni, C. A. Klebanoff, D. C. Palmer, C. Wrzesinski, K. Kerstann, Z. Yu, S. E. Finkelstein, M. R. Theoret, S. A. Rosenberg, N. P. Restifo, Acquisition of full effector function in vitro paradoxically impairs the in vivo antitumor efficacy of adoptively transferred CD8+ T cells. J Clin Invest 115, 1616–1626 (2005).

7. A. Chow, K. Perica, C. A. Klebanoff, J. D. Wolchok, Clinical implications of T cell exhaustion for cancer immunotherapy. Nat Rev Clin Oncol 19, 775–790 (2022).

8. M. Shourian, J. C. Beltra, B. Bourdin, H. Decaluwe, Common gamma chain cytokines and CD8 T cells in cancer. Semin Immunol 42, 101307 (2019).

9. J. C. Beltra, H. Decaluwe, Cytokines and persistent viral infections. Cytokine 82, 4–15 (2016).

10. W. J. Leonard, J. X. Lin, J. J. O’Shea, The gamma(c) Family of Cytokines: Basic Biology to Therapeutic Ramifications. Immunity 50, 832–850 (2019).

11. V. Kalia, S. Sarkar, S. Subramaniam, W. N. Haining, K. A. Smith, R. Ahmed, Prolonged interleukin-2Ralpha expression on virus-specific CD8+ T cells favors terminal-effector differentiation in vivo. Immunity 32, 91–103 (2010).

12. J. C. Markley, M. Sadelain, IL-7 and IL-21 are superior to IL-2 and IL-15 in promoting human T cell-mediated rejection of systemic lymphoma in immunodeficient mice. Blood 115, 3508–3519 (2010).

13. C. S. Hinrichs, R. Spolski, C. M. Paulos, L. Gattinoni, K. W. Kerstann, D. C. Palmer, C. A. Klebanoff, S. A. Rosenberg, W. J. Leonard, N. P. Restifo, IL-2 and IL-21 confer opposing differentiation programs to CD8+ T cells for adoptive immunotherapy. Blood 111, 5326–5333 (2008).

14. R. Zeng, R. Spolski, S. E. Finkelstein, S. Oh, P. E. Kovanen, C. S. Hinrichs, C. A. Pise-Masison, M. F. Radonovich, J. N. Brady, N. P. Restifo, J. A. Berzofsky, W. J. Leonard, Synergy of IL-21 and IL-15 in regulating CD8+ T cell expansion and function. J Exp Med 201, 139–148 (2005).

15. N. Cieri, B. Camisa, F. Cocchiarella, M. Forcato, G. Oliveira, E. Provasi, A. Bondanza, C. Bordignon, J. Peccatori, F. Ciceri, M. T. Lupo-Stanghellini, F. Mavilio, A. Mondino, S. Bicciato, A. Recchia, C. Bonini, IL-7 and IL-15 instruct the generation of human memory stem T cells from naive precursors. Blood 121, 573–584 (2013).

16. A. M. Ring, J. X. Lin, D. Feng, S. Mitra, M. Rickert, G. R. Bowman, V. S. Pande, P. Li, I. Moraga, R. Spolski, E. Ozkan, W. J. Leonard, K. C. Garcia, Mechanistic and structural insight into the functional dichotomy between IL-2 and IL-15. Nat Immunol, (2012).

17. T. A. Waldmann, The shared and contrasting roles of IL2 and IL15 in the life and death of normal and neoplastic lymphocytes: implications for cancer therapy. Cancer Immunol Res 3, 219–227 (2015).

18. J. C. Beltra, S. Bourbonnais, N. Bedard, T. Charpentier, M. Boulange, E. Michaud, I. Boufaied, J. Bruneau, N. H. Shoukry, A. Lamarre, H. Decaluwe, IL2Rbeta-dependent signals drive terminal exhaustion and suppress memory development during chronic viral infection. Proc Natl Acad Sci U S A 113, E5444–5453 (2016).

19. B. C. Miller, D. R. Sen, R. Al Abosy, K. Bi, Y. V. Virkud, M. W. LaFleur, K. B. Yates, A. Lako, K. Felt, G. S. Naik, M. Manos, E. Gjini, J. R. Kuchroo, J. J. Ishizuka, J. L. Collier, G. K. Griffin, S. Maleri, D. E. Comstock, S. A. Weiss, F. D. Brown, A. Panda, M. D. Zimmer, R. T. Manguso, F. S. Hodi, S. J. Rodig, A. H. Sharpe, W. N. Haining, Subsets of exhausted CD8(+) T cells differentially mediate tumor control and respond to checkpoint blockade. Nat Immunol 20, 326–336 (2019).

20. L. M. McLane, M. S. Abdel-Hakeem, E. J. Wherry, CD8 T Cell Exhaustion During Chronic Viral Infection and Cancer. Annu Rev Immunol 37, 457–495 (2019).

21. W. H. Hudson, J. Gensheimer, M. Hashimoto, A. Wieland, R. M. Valanparambil, P. Li, J. X. Lin, B. T. Konieczny, S. J. Im, G. J. Freeman, W. J. Leonard, H. T. Kissick, R. Ahmed, Proliferating Transitory T Cells with an Effector-like Transcriptional Signature Emerge from PD-1(+) Stem-like CD8(+) T Cells during Chronic Infection. Immunity 51, 1043–1058 e1044 (2019).

22. R. Zander, D. Schauder, G. Xin, C. Nguyen, X. Wu, A. Zajac, W. Cui, CD4(+) T Cell Help Is Required for the Formation of a Cytolytic CD8(+) T Cell Subset that Protects against Chronic Infection and Cancer. Immunity 51, 1028–1042 e1024 (2019).

23. J. C. Beltra, S. Manne, M. S. Abdel-Hakeem, M. Kurachi, J. R. Giles, Z. Chen, V. Casella, S. F. Ngiow, O. Khan, Y. J. Huang, P. Yan, K. Nzingha, W. Xu, R. K. Amaravadi, X. Xu, G. C. Karakousis, T. C. Mitchell, L. M. Schuchter, A. C. Huang, E. J. Wherry, Developmental Relationships of Four Exhausted CD8(+) T Cell Subsets Reveals Underlying Transcriptional and Epigenetic Landscape Control Mechanisms. Immunity 52, 825–841 e828 (2020).

24. B. Bengsch, T. Ohtani, O. Khan, M. Setty, S. Manne, S. O’Brien, P. F. Gherardini, R. S. Herati, A. C. Huang, K. M. Chang, E. W. Newell, N. Bovenschen, D. Pe’er, S. M. Albelda, E. J. Wherry, Epigenomic-Guided Mass Cytometry Profiling Reveals Disease-Specific Features of Exhausted CD8 T Cells. Immunity 48, 1029–1045 e1025 (2018).

25. T. A. Waldmann, The biology of interleukin-2 and interleukin-15: implications for cancer therapy and vaccine design. Nat Rev Immunol 6, 595–601 (2006).

26. M. M. Staron, S. M. Gray, H. D. Marshall, I. A. Parish, J. H. Chen, C. J. Perry, G. Cui, M. O. Li, S. M. Kaech, The transcription factor FoxO1 sustains expression of the inhibitory receptor PD-1 and survival of antiviral CD8(+) T cells during chronic infection. Immunity 41, 802–814 (2014).

27. Y. Liu, N. Zhou, L. Zhou, J. Wang, Y. Zhou, T. Zhang, Y. Fang, J. Deng, Y. Gao, X. Liang, J. Lv, Z. Wang, J. Xie, Y. Xue, H. Zhang, J. Ma, K. Tang, Y. Fang, F. Cheng, C. Zhang, B. Dong, Y. Zhao, P. Yuan, Q. Gao, H. Zhang, F. Xiao-Feng Qin, B. Huang, IL-2 regulates tumor-reactive CD8(+) T cell exhaustion by activating the aryl hydrocarbon receptor. Nat Immunol 22, 358–369 (2021).

28. F. Mo, Z. Yu, P. Li, J. Oh, R. Spolski, L. Zhao, C. R. Glassman, T. N. Yamamoto, Y. Chen, F. M. Golebiowski, D. Hermans, S. Majri-Morrison, L. K. Picton, W. Liao, M. Ren, X. Zhuang, S. Mitra, J. X. Lin, L. Gattinoni, J. D. Powell, N. P. Restifo, K. C. Garcia, W. J. Leonard, An engineered IL-2 partial agonist promotes CD8(+) T cell stemness. Nature 597, 544–548 (2021).

29. L. Gattinoni, D. E. Speiser, M. Lichterfeld, C. Bonini, T memory stem cells in health and disease. Nat Med 23, 18–27 (2017).

30. E. J. Wherry, M. Kurachi, Molecular and cellular insights into T cell exhaustion. Nat Rev Immunol 15, 486–499 (2015).

31. D. T. Utzschneider, M. Charmoy, V. Chennupati, L. Pousse, D. P. Ferreira, S. Calderon-Copete, M. Danilo, F. Alfei, M. Hofmann, D. Wieland, S. Pradervand, R. Thimme, D. Zehn, W. Held, T Cell Factor 1-Expressing Memory-like CD8(+) T Cells Sustain the Immune Response to Chronic Viral Infections. Immunity 45, 415–427 (2016).

32. J. E. Wu, S. Manne, S. F. Ngiow, A. E. Baxter, H. Huang, E. Freilich, M. L. Clark, J. H. Lee, Z. Chen, O. Khan, R. P. Staupe, Y. J. Huang, J. Shi, J. R. Giles, E. J. Wherry, In vitro modeling of CD8(+) T cell exhaustion enables CRISPR screening to reveal a role for BHLHE40. Sci Immunol 8, eade3369 (2023).

33. M. Zhao, C. H. Kiernan, C. J. Stairiker, J. L. Hope, L. G. Leon, M. van Meurs, I. Brouwers-Haspels, R. Boers, J. Boers, J. Gribnau, I. W. F. J. van, E. M. Bindels, R. M. Hoogenboezem, S. J. Erkeland, Y. M. Mueller, P. D. Katsikis, Rapid in vitro generation of bona fide exhausted CD8+ T cells is accompanied by Tcf7 promotor methylation. PLoS Pathog 16, e1008555 (2020).

34. E. W. Weber, K. R. Parker, E. Sotillo, R. C. Lynn, H. Anbunathan, J. Lattin, Z. Good, J. A. Belk, B. Daniel, D. Klysz, M. Malipatlolla, P. Xu, M. Bashti, S. Heitzeneder, L. Labanieh, P. Vandris, R. G. Majzner, Y. Qi, K. Sandor, L. C. Chen, S. Prabhu, A. J. Gentles, T. J. Wandless, A. T. Satpathy, H. Y. Chang, C. L. Mackall, Transient rest restores functionality in exhausted CAR-T cells through epigenetic remodeling. Science 372, (2021).

35. A. B. L. Colamartino, P. Bifsha, C. Colas, C. Tremblay-Laganière, S. Nicoletti, M. Guiot, A. Boldici, Y. Li, E. Haddad, A New Specific Promoter Allow Hematopoietic Stem Cell Immunotherapy Approach Against Acute Lymphoblastic Leukemia Using Chimeric Antigen Receptor. Biology of Blood and Marrow Transplantation, (2019).

36. A. B. L. Colamartino, W. Lemieux, P. Bifsha, S. Nicoletti, N. Chakravarti, J. Sanz, H. Romero, S. Selleri, K. Beland, M. Guiot, C. Tremblay-Laganiere, R. Dicaire, L. Barreiro, D. A. Lee, E. Verhoeyen, E. Haddad, Efficient and Robust NK-Cell Transduction With Baboon Envelope Pseudotyped Lentivector. Front Immunol 10, 2873 (2019).

37. W. Haso, D. W. Lee, N. N. Shah, M. Stetler-Stevenson, C. M. Yuan, I. H. Pastan, D. S. Dimitrov, R. A. Morgan, D. J. FitzGerald, D. M. Barrett, A. S. Wayne, C. L. Mackall, R. J. Orentas, Anti-CD22-chimeric antigen receptors targeting B-cell precursor acute lymphoblastic leukemia. Blood 121, 1165–1174 (2013).

38. M. Hashimoto, S. J. Im, K. Araki, R. Ahmed, Cytokine-Mediated Regulation of CD8 T-Cell Responses During Acute and Chronic Viral Infection. Cold Spring Harb Perspect Biol 11, (2019).

39. D. G. Brooks, A. M. Lee, H. Elsaesser, D. B. McGavern, M. B. Oldstone, IL-10 blockade facilitates DNA vaccine-induced T cell responses and enhances clearance of persistent virus infection. J Exp Med 205, 533–541 (2008).

40. F. Bertrand, J. Rochotte, C. Colacios, A. Montfort, A. F. Tilkin-Mariame, C. Touriol, P. Rochaix, I. Lajoie-Mazenc, N. Andrieu-Abadie, T. Levade, H. Benoist, B. Segui, Blocking Tumor Necrosis Factor alpha Enhances CD8 T-cell-Dependent Immunity in Experimental Melanoma. Cancer Res 75, 2619–2628 (2015).

41. M. A. Huseni, L. Wang, J. E. Klementowicz, K. Yuen, B. Breart, C. Orr, L. F. Liu, Y. Li, V. Gupta, C. Li, D. Rishipathak, J. Peng, Y. Senbabaoglu, Z. Modrusan, S. Keerthivasan, S. Madireddi, Y. J. Chen, E. J. Fraser, N. Leng, H. Hamidi, H. Koeppen, J. Ziai, K. Hashimoto, M. Fasso, P. Williams, D. F. McDermott, J. E. Rosenberg, T. Powles, L. A. Emens, P. S. Hegde, I. Mellman, S. J. Turley, M. S. Wilson, S. Mariathasan, L. Molinero, M. Merchant, N. R. West, CD8(+) T cell-intrinsic IL-6 signaling promotes resistance to anti-PD-L1 immunotherapy. Cell Rep Med 4, 100878 (2023).

42. Y. Rochman, R. Spolski, W. J. Leonard, New insights into the regulation of T cells by gamma(c) family cytokines. Nat Rev Immunol 9, 480–490 (2009).

43. S. Ghaffari, M. Torabi-Rahvar, S. Aghayan, Z. Jabbarpour, K. Moradzadeh, A. Omidkhoda, N. Ahmadbeigi, Optimizing interleukin-2 concentration, seeding density and bead-to-cell ratio of T-cell expansion for adoptive immunotherapy. BMC Immunol 22, 43 (2021).

44. B. Kwon, The two faces of IL-2: a key driver of CD8(+) T-cell exhaustion. Cell Mol Immunol 18, 1641–1643 (2021).

45. M. Felices, A. J. Lenvik, R. McElmurry, S. Chu, P. Hinderlie, L. Bendzick, M. A. Geller, J. Tolar, B. R. Blazar, J. S. Miller, Continuous treatment with IL-15 exhausts human NK cells via a metabolic defect. JCI Insight 3, (2018).

46. K. Mueller, O. Schweier, H. Pircher, Efficacy of IL-2-versus IL-15-stimulated CD8 T cells in adoptive immunotherapy. Eur J Immunol 38, 2874–2885 (2008).

47. J. C. Beltra, M. S. Abdel-Hakeem, S. Manne, Z. Zhang, H. Huang, M. Kurachi, L. Su, L. Picton, S. F. Ngiow, Y. Muroyama, V. Casella, Y. J. Huang, J. R. Giles, D. Mathew, J. Belman, M. Klapholz, H. Decaluwe, A. C. Huang, S. L. Berger, K. C. Garcia, E. J. Wherry, Stat5 opposes the transcription factor Tox and rewires exhausted CD8(+) T cells toward durable effector-like states during chronic antigen exposure. Immunity 56, 2699–2718 e2611 (2023).

48. L. Gattinoni, E. Lugli, Y. Ji, Z. Pos, C. M. Paulos, M. F. Quigley, J. R. Almeida, E. Gostick, Z. Yu, C. Carpenito, E. Wang, D. C. Douek, D. A. Price, C. H. June, F. M. Marincola, M. Roederer, N. P. Restifo, A human memory T cell subset with stem cell-like properties. Nat Med 17, 1290–1297 (2011).

49. D. Zehn, R. Thimme, E. Lugli, G. P. de Almeida, A. Oxenius, ’Stem-like’ precursors are the fount to sustain persistent CD8(+) T cell responses. Nat Immunol 23, 836–847 (2022).

50. T. Gebhardt, S. L. Park, I. A. Parish, Stem-like exhausted and memory CD8(+) T cells in cancer. Nat Rev Cancer 23, 780–798 (2023).

51. J. S. Bowers, K. Majchrzak, M. H. Nelson, B. A. Aksoy, M. M. Wyatt, A. S. Smith, S. R. Bailey, L. R. Neal, J. E. Hammerbacher, C. M. Paulos, PI3Kdelta Inhibition Enhances the Antitumor Fitness of Adoptively Transferred CD8(+) T Cells. Front Immunol 8, 1221 (2017).

52. C. A. Klebanoff, J. G. Crompton, A. J. Leonardi, T. N. Yamamoto, S. S. Chandran, R. L. Eil, M. Sukumar, S. K. Vodnala, J. Hu, Y. Ji, D. Clever, M. A. Black, D. Gurusamy, M. J. Kruhlak, P. Jin, D. F. Stroncek, L. Gattinoni, S. A. Feldman, N. P. Restifo, Inhibition of AKT signaling uncouples T cell differentiation from expansion for receptor-engineered adoptive immunotherapy. JCI Insight 2, (2017).

53. R. R. Rao, Q. Li, K. Odunsi, P. A. Shrikant, The mTOR kinase determines effector versus memory CD8+ T cell fate by regulating the expression of transcription factors T-bet and Eomesodermin. Immunity 32, 67–78 (2010).

54. C. Mathieu, J. C. Beltra, T. Charpentier, S. Bourbonnais, J. P. Di Santo, A. Lamarre, H. Decaluwe, IL-2 and IL-15 regulate CD8+ memory T-cell differentiation but are dispensable for protective recall responses. Eur J Immunol 45, 3324–3338 (2015).

55. K. Speer, A. Poole, M. Rodriguez-Lanetty, RNA extraction from adult Aiptasia. Protocols.io, (2018).

56. K. E. Pauken, M. A. Sammons, P. M. Odorizzi, S. Manne, J. Godec, O. Khan, A. M. Drake, Z. Chen, D. R. Sen, M. Kurachi, R. A. Barnitz, C. Bartman, B. Bengsch, A. C. Huang, J. M. Schenkel, G. Vahedi, W. N. Haining, S. L. Berger, E. J. Wherry, Epigenetic stability of exhausted T cells limits durability of reinvigoration by PD-1 blockade. Science 354, 1160–1165 (2016).

57. D. R. Sen, J. Kaminski, R. A. Barnitz, M. Kurachi, U. Gerdemann, K. B. Yates, H. W. Tsao, J. Godec, M. W. LaFleur, F. D. Brown, P. Tonnerre, R. T. Chung, D. C. Tully, T. M. Allen, N. Frahm, G. M. Lauer, E. J. Wherry, N. Yosef, W. N. Haining, The epigenetic landscape of T cell exhaustion. Science 354, 1165–1169 (2016).

