## Supplemental material for "Inhibition of IL2Rß-dependent STAT5 activity supports T-cell stemness and augments antitumor efficacy of CD8^+^ T cells by preventing T-cell exhaustion"

### Supplementary Material

#### Supplemental Figures

**Figure S1.** IL2Rβ deficiency maintains CD8+ T cells in a progenitor exhausted state during LCMV Cl.13 infection.

**Figure S2.** Inhibition of STAT5 increases stemness of *in vitro* stimulated $CD8^+$ T cells.

**Figure S3.** Inhibition of JAK molecules with tofacitinib or JAK3i favors the development of stem-like progenitors *in vitro*.

#### Supplemental Tables (excel files)

**Table S1.** Differential gene expression between IL2Rβ high and IL2Rβ low $CD8^+$ $tetramer^+$ $CD11a^+$ T cells at day 21 post-LCMV Cl.13 infection.

**Table S2.** Differential gene expression of *in vitro* cultured $CD8^+$ T cells stimulated with continuous IL-2 and peptide versus naive $CD8^+$ T cells.

**Table S3.** Differential gene expression of *in vitro* cultured $CD8^+$ T cells stimulated with continuous IL-15 and peptide versus naive $CD8^+$ T cells.

**Table S4.** Differential gene expression of *in vitro* cultured $CD8^+$ T cells stimulated with IL-2 and STAT5i (IL-2S5i) versus IL-2.

**Table S5.** Differential gene expression of *in vitro* cultured $CD8^+$ T cells stimulated with IL-15 and STAT5i (IL-15S5i) versus IL-15.

**Fig. S1. IL2Rβ deficiency maintains CD8$^+$ T cells in a progenitor exhausted state during LCMV Cl.13 infection.**

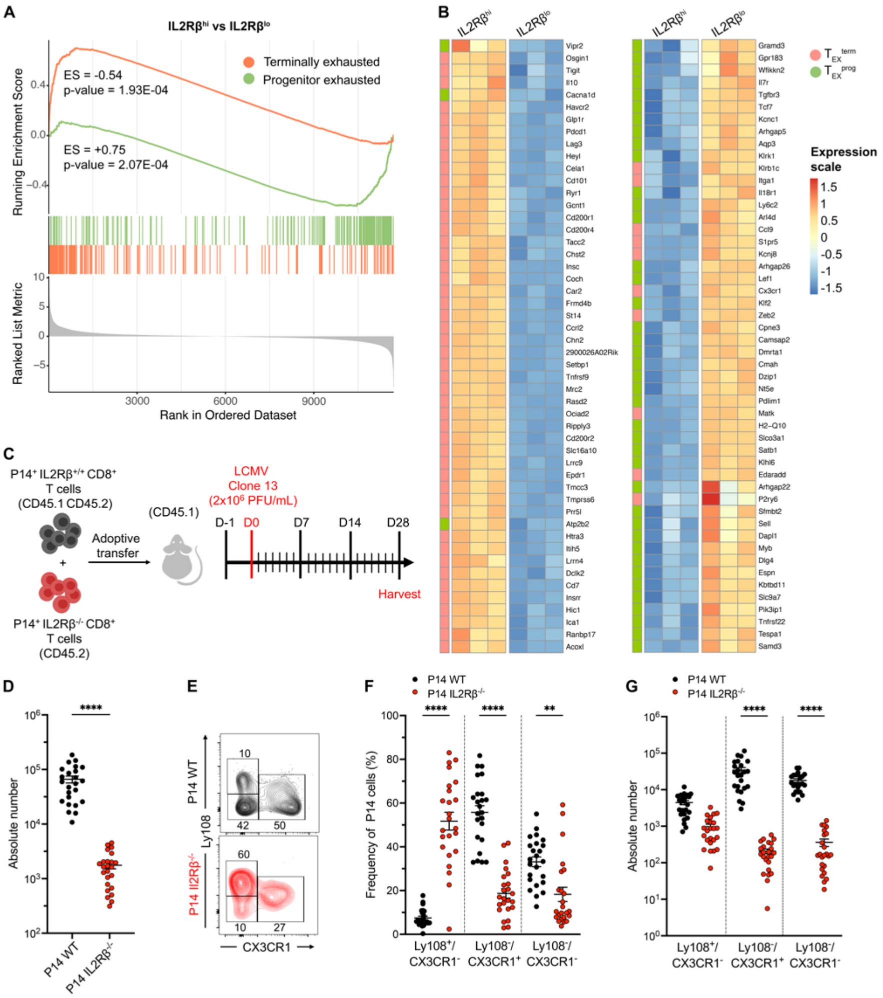


**(A)** Pre-ranked GSEA of DE gene profiles comparing CD8$^+$ Tet$^+$ CD11a$^+$ IL2Rβ$^{hi}$ versus IL2Rβ$^{lo}$ T cells at day 21 post-LCMV Cl.13 infection to Miller *et al.*'s gene signatures associated with progenitor (Slamf6$^+$Tim-3$^-$, green line) and terminally exhausted (Slamf6$^+$Tim-3$^-$, orange line) (GSE123235) (*52*). ES: enrichment score. **(B)** Heatmap displaying scaled gene expression data of individual IL2Rβ$^{hi}$ or IL2Rβ$^{lo}$ replicates (n=3) for the top 50 up/down regulated genes associated with progenitor and terminally exhausted CD8$^+$ T cells from GSE123235 dataset ($\log_2$ FC $\geq 0.5$ and FDR < 5%). **(C)** Schematic of the experimental design for adoptive co-transfer studies. P14$^+$ IL-2Rβ$^{+/+}$ (P14 WT, black) (CD45.1 CD45.2) and P14$^+$ IL2Rβ$^{-/-}$ (red) (CD45.2) T cells were co-transferred into B6 mice (CD45.1$^+$) 24 hours prior to infection with LCMV Cl.13. Cells were analyzed at day +28 p.i.. **(D)** Scatter plot of the absolute number of $P14^{+}$ WT (black circles) and $P14^{+}$ $IL2R\beta^{-/-}$ (red circles) T cells 28 days p.i. (**E**) Representative flow cytometry plots for Ly108 and CX3CR1 co-expression. Scatter plots of **(F)** the frequency and **(G)** absolute number of $Ly108^{+}CX3CR1^{-}$, $Ly108^{-}CX3CR1^{+}$ and $Ly108^{-}CX3CR1^{-}$ cell populations. Data are combined (D, F and G) from two independent experiments (total of 24 mice). Error bars indicate means ± SEM. *$P < 0.05$; **$P < 0.01$; ***$P < 0.001$; ****$P < 0.0001$, using t-test (D) or one-way ANOVA (F, G).

**Fig. S2. Inhibition of STAT5 increases stemness of *in vitro* stimulated $CD8^+$ T cells.**

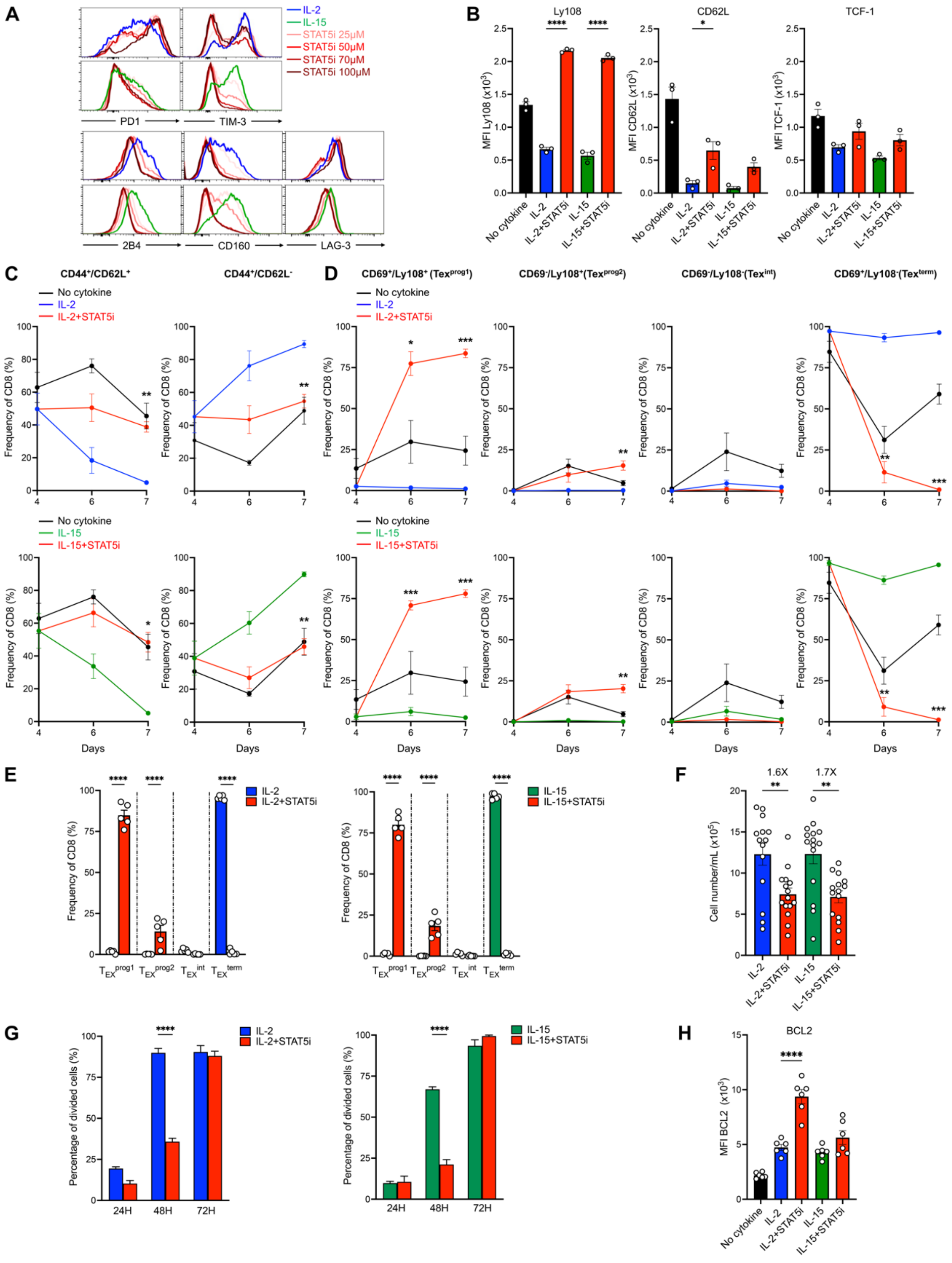


P14 $CD8^+$ T cells were expanded *in vitro* with gp33-pulsed DCs alone (no cytokine), or combined with IL-2 (100 ng/mL, blue) or IL-15 (200 ng/mL, green) in the presence or absence of STAT5i

(100 μM unless otherwise indicated, red), as described in **Fig. 3B**. (**A**) Histogram of indicated IRs at different doses of STAT5i. (**B**) Bar graphs of the MFI of Ly108, CD62L and TCF-1 at day 7. (**C**) Frequency of $CD44^+CD62L^+$ and $CD44^+CD62L^-$ populations at 4, 6 and 7 days of culture with or without STAT5i and IL-2 or IL-15 stimulation. **(D, E)** Frequency of $T_{EX}^{prog1}$ ($CD69^+Ly108^+$), $T_{EX}^{prog2}$ ($CD69^-Ly108^+$), T $T_{EX}^{int}$ ($CD69^-Ly108^-$), and $T_{EX}^{term}$ ($CD69^+Ly108^-$) populations at day 4, 6 and 7, in IL-2 **(D)** or IL-15 **(E)** +/- STAT5i culture conditions. **(F**) Bar graphs of the absolute counts of $CD8^+$ T cells at day 7. **(G)** P14 $CD8^+$ T cells were stained with CFSE on day 4 of the culture. Bar graph of the percentage of divided cells after 24, 48, and 72 hours. **(H)** Bar graphs of the MFI of BCL2 day 7. Data is representative of at least three independent experiments (B, G, H), or are combined from 3-5 independent experiments (C-F). Error bars indicate means ± SEM. *$P < 0.05$; **$P < 0.01$; ***$P < 0.001$; ****$P < 0.0001$, using one-way (B, E-H) or two-way ANOVA (C and D).

**Fig. S3. Inhibition of JAK molecules with tofacitinib or JAK3i favors the development of stem-like progenitors *in vitro*.**

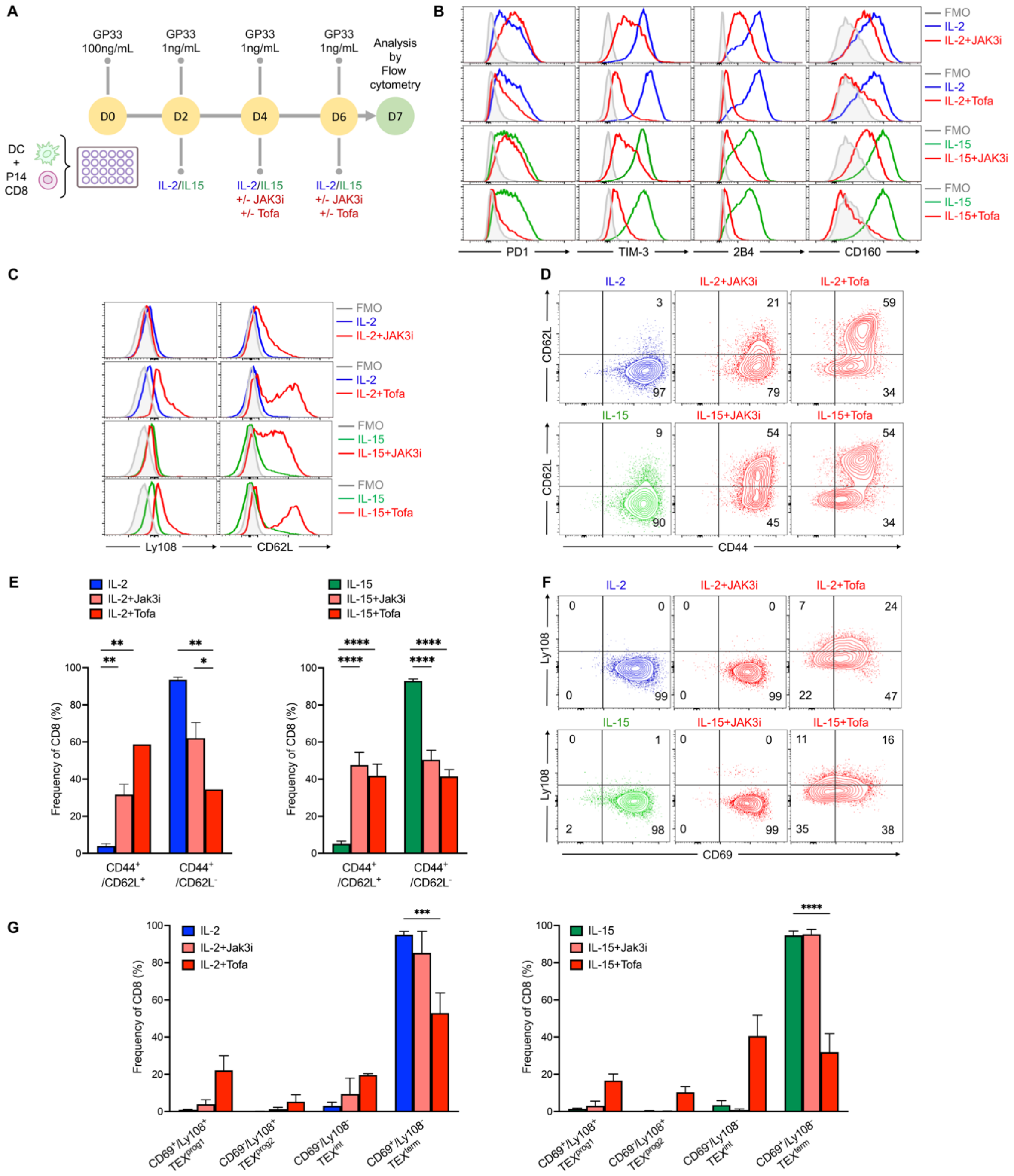

**(A)** P14 CD8$^{+}$ T cells were expanded *in vitro* with gp33-pulsed DCs and IL-2 (100 ng/mL, blue) or IL-15 (200 ng/mL, green) as described in **Fig. 2A**. Tofacitinib (Tofa, 1μM, dark red) or JAK3 inhibitor (Jak3i, 3μM, light red) were added to the culture on days 4 and 6. Cells were analyzed at day 7. (**B-C**) Histograms of the indicated molecules. (**D, F**) Contour plots and of (**D**) CD44/CD62L and (**F**) CD69/Ly108 co-expression. (**E**) Bar graphs of the frequency of CD44$^{+}$CD62L$^{+}$ and CD44$^{+}$CD62L$^{-}$ cells at day 7. (G) Bar graphs of the frequency of $T_{EX}^{prog1}$ (CD69$^{+}$Ly108$^{+}$), $T_{EX}^{prog2}$ (CD69$^{-}$Ly108$^{+}$), $T_{EX}^{int}$ (CD69$^{-}$Ly108$^{-}$), and $T_{EX}^{term}$ (CD69$^{+}$Ly108$^{-}$) populations at day 7. Data are representative (B, C, D and F) or combined (E and G) from two independent experiments. Error bars indicate means ± SEM. *P < 0.05; **P < 0.01; ***P < 0.001; ****P < 0.0001, using one-Way ANOVA.
